# Inducing a synergistic anti-obesity effect by increasing the bioavailability of the flavonoid rutin with a *L. plantarum* species

**DOI:** 10.1101/2024.10.28.620556

**Authors:** Youn-Goo Kang, Seongjae Jang, Jongcheol Seo, Ah-Ram Kim

## Abstract

Flavonoids, plant-derived compounds, are broadly categorized into glycoside (sugar-bonded) and aglycone (sugar-removed) forms and are predominantly found as glycosides in nature. Rutin, a glycoside, is a widely recognized flavonoid with significant potential as an anti-obesity agent. However, its low bioavailability in the body presents challenges in obesity treatment. We aimed to enhance the bioavailability and anti-obesity effect of rutin through microbial involvement. *Lactiplantibacillus plantarum* HAC03, one of the candidates, demonstrated the ability to hydrolyze rutin into isoquercetin and quercetin *in vitro*. In a diet-induced obesity mouse model, the combination of rutin and the *L. plantarum* strain resulted in significant weight loss, reduced adipocyte size and lowered obesity-related biomarkers in the blood, decreased fat-synthesis related gene expression, and increased fatty acid β-oxidation related gene expression compared to other test groups. This includes groups treated with rutin or quercetin alone or in combination with a different species from the same *Lactobacillus* genus, known for its anti-obesity effect but lacking the ability to hydrolyze rutin. This synergistic combination also alleviated insulin resistance and reduced fat in the liver. Gut microbiota analysis revealed localization of *L. plantarum* in the ileum and beneficial changes in disrupted microbiota in the intestine. These findings provide insights into underlying mechanisms causing the synergistic effect and suggest a novel combination that is as safe as microbial monotherapies with *L. plantarum* or *L. rhamnosus* but more effective in anti-obesity treatment.

## Introduction

Obesity is a chronic condition characterized by abnormal or excessive fat accumulation, which poses health risks such as hypertension, type II diabetes, dyslipidemia, and fatty liver disease.^1,2^ Body mass index (BMI) is used to classify overweight and obesity, with a BMI over 25 kg/m^2^ considered overweight and over 30 kg/m^2^ considered obese.^3^ According to the World Obesity Atlas 2023 presented by the World Obesity Federation, more than 2.6 billion people (38% of the global population) were classified as overweight or obese in 2020. This number is expected to increase to over 4 billion people (50% of the global population) by 2035.^4^

Natural compounds in plant extracts have been used in traditional folk medicine for weight management and metabolic disorders.^5^ Flavonoids, one of the phytochemicals, are found in over 10,000 different varieties and have been discovered in various plants, including vegetables and fruits.^6^ Rutin is one of the most abundant flavonoids in plants and has been shown significant potential as an anti-obesity agent.^7,8^ However, limited bioavailability has hindered its widespread use an anti-obesity treatments.^9^

Flavonoids share a common carbon skeleton structure, consisting of two phenolic rings and a heterocyclic ring in a C_6_-C_3_-C_6_ configuration.^10^ They can be broadly classified into aglycones and glycosides (the latter referring to aglycones with sugar molecules attached), which differ in their absorption mechanisms and bioavailability in the body.^11–13^ Due to their higher hydrophobicity compared to their glycoside forms, aglycones are generally more readily absorbed in the intestine. For example, quercetin, an aglycone and a hydrolyzed derivative of rutin, is well absorbed through passive diffusion, particularly in the small intestine.^14–16^ However, most glycosides such as rutin cannot be efficiently absorbed by the human body since the human intestine lacks the necessary enzymes for their hydrolysis.^17^ Consequently, most glycosides rely on the assistance of gut microbiota in the large intestine for absorption.^18^

In the case of rutin, it is hydrolyzed by microbial enzymes such as α-L-rhamnosidase and β-D-glucosidase produced by gut microbiota, including *Lactobacillus* spp., *Bacteroides* spp., and *Enterococcus* spp.^19^ α-L-rhamnosidase converts rutin into isoquercetin and β-D-glucosidase converts isoquercetin into quercetin, which can undergoes further conversion into phenolic acids in the intestine, facilitating its absorption into colon enterocytes.^20,21^ However, certain glycosides, such as isoquercetin, have the capability to be directly absorbed in the human small intestine. Typically, these special glycosides are associated with glucose.^22^ Isoquercetin absorption in the small intestine can occur via two pathways; it can either be absorbed into enterocytes after enzymatic hydrolysis to quercetin by lactase phlorizin hydrolase (LPH) or be directly transported to enterocytes through sodium-glucose cotransporter 1 (SGLT-1) and then undergo hydrolysis into quercetin by cytosolic β-glucosidase (CBG) within the enterocytes.^16^ In pharmacokinetic studies, isoquercetin demonstrated the highest bioavailability, followed by quercetin (second highest), whereas rutin exhibited the lowest bioavailability.^23^ Both of these rutin derivatives, like rutin itself, exhibited an anti-obesity effect when administered individually.^24^

This study aimed to enhance the bioavailability of rutin with the help of a beneficial bacterium capable of hydrolyzing rutin and ultimately increase its anti-obesity effect. We selected two testing species from the same *Lactobacillus* genus — *Lactiplantibacillus plantarum* (formerly known as *Lactobacillus plantarum*) and *Lactobacillus rhamnosus*. While both share proven anti-obesity properties, they differ in their ability to hydrolyze rutin. In our *in vitro* experiments, *L. plantarum* HAC03, isolated from Korean traditional white kimchi demonstrated strong rutin hydrolysis capability. In contrast, *L. rhamnosus* ATCC 53103 did not hydrolyze rutin. In a diet-induced obesity (DIO) mouse model, we found that the combination of rutin and the *L. plantarum* strain induces a synergistic anti-obesity effect compared to other test groups including the combination of rutin and the *L. rhamnosus* strain. Based on *in vitro* and *in vivo* analyses, we propose the molecular mechanisms that underlie the synergy between rutin and *L. plantarum*.

## Results

### *L. plantarum* HAC03 can hydrolyze rutin into isoquercetin and quercetin

In selecting *Lactobacillus* candidates for testing, we referred to the KEGG Orthology (KO) database and relevant literature. From 68 *Lactobacillus* spp. listed in the KO, 25 strains possessed α-L-rhamnosidase (EC 3.2.1.40) encoded by *RamA* (K05989), 22 strains had β-D-glucosidase (EC 3.2.1.21) encoded by *bglX* (K05349), and 65 strains possessed phospho-β-D-glucosidase (EC 3.2.1.86) encoded by *bglA* (K01223) (Supplementary Table 1). However, it appears that only three *Lactobacillus* strains — *L. acidophilus, L. plantarum and L. rhamnosus —* have all the necessary enzymes to convert rutin into quercetin. According to relevant literature, among these three, *L. plantarum* exhibited the most outstanding ability to hydrolyze rutin, while *L. rhamnosus*, contrary to information in the KO database, was reported to lack the ability.^25^ Our *in vitro* experiment results also aligned with these findings. Given that *L. plantarum* and *L. rhamnosus* both exhibit proven anti-obesity properties but significantly differ in their ability to hydrolyze rutin, we selected them for comparative experiments.

In order to assess the rutin hydrolysis ability of the candidate strains, we first established specific growth conditions for the 6 day experimentation of the candidate strain: 1) CaCO_3_ was introduced into the culture medium as a pH-regulating agent to prevent a rapid decrease in pH during strain growth.^26^ 2) a glucose-free MRS medium was employed to augment the rutin hydrolysis ability of the candidates.^27^ All experimental groups tested in this study used a MRS medium with no glucose and CaCO_3_ added, named MRS-g+C.

Rutin and its derivatives in the MRS medium, without glucose and CaCO_3_ additives, were measured using High-Performance Liquid Chromatography with diode-array detector and mass spectrometry (HPLC-DAD-MS) under various culture conditions. In the negative control (MRS-g+C only), rutin and its derivatives—isoquercetin and quercetin were not detected after six days of incubation. When the MRS-g+C medium was cultured solely with rutin for six days, rutin alone was detected. However, in the medium with rutin and *L. plantarum* HAC03, all three compounds—rutin, isoquercetin, and quercetin—were detected (Figure 2a). In the medium containing rutin and the *L. plantarum* strain, the gradual decrease in rutin concentration over six days, accompanied by an increase in isoquercetin and quercetin concentrations, demonstrates that the *L. plantarum* strain acts as the biocatalyst for breaking down rutin into its derivatives (Figure 2d). In contrast, in the medium with rutin and *L. rhamnosus* ATCC 53103, rutin derivatives were not detected while the concentration of rutin was slightly decreased over time (Supplementary Figure 1a and Supplementary Figure 1d). We used this *L. rhamnosus* strain that lacks the ability to hydrolyze rutin but is well known for strong anti-obesity effect ^28,29^ as a positive control.

### Combination of rutin and *L. plantarum* HAC03 has a synergic effect on losing body weight

To investigate the anti-obesity effect of the combination of rutin and *L. plantarum* HAC03, we administered 1) rutin or 2) quercetin (the end product of rutin hydrolysis by *L. plantarum* HAC03) or 3) the two bacterial strains (*L. plantarum* HAC03 and *L. rhamnosus* ATCC 53103) individually or in combination to a diet induced obesity (DIO) mouse model for a period of 14 weeks (Figure 1a). After 14 weeks of administration, all test groups, including the low-fat diet group (LFD), had significantly reduced body weight gain compared to those in the high-fat diet group (HFD) (Figure 3a and b). The rutin treatment group (R) exhibited significantly lower body weight gain than that in the quercetin treatment group (Q) (p-value = 0.017) (Figure 3a and b). The combination of rutin and *L. plantarum* HAC03 treatment group (R+LP) showed the least body weight gain (p-value < 0.001) compared to the HFD group (Figure 3b). Particularly, the R+LP group demonstrated a more significant reduction in body weight compared to that in the *L. plantarum* HAC03 treatment alone (LP) or combination of quercetin and *L. plantarum* HAC03 (Q+LP) (p-value compared with LP = 0.042, p-value compared with Q+LP = 0.008) (Figure 3b). On the contrary, whether administered alone or in combination with the testing flavonoids, *L. rhamnosus* ATCC 53103 did not exhibit significant differences in body weight gain among the groups (p-value between LR and Q+LR = 0.273, p-value between LR and R+LR = 0.090) (Figure 3b). However, when comparing the average daily food intake over 14 weeks, there was no difference observed among the groups fed HFD, except for the LFD group (Supplementary Figure 2a). With respect to the weight of the liver and four adipose tissues, the R+LP group exhibited a significant reduction across all measured tissues when compared to the HFD group (p-value < 0.001) (Figure 3c). Particularly, among the four adipose tissues, the greatest statistical difference was observed in mesenteric adipose tissue (MAT) when compared to that in the HFD group (p-value = 0.000016) (Figure 3c). Measurements of obesity-related biochemical markers (triglyceride, total cholesterol, low-density lipoprotein (LDL), and high-density lipoprotein (HDL)) in the blood serum revealed that the R+LP group exhibited the most significant reduction compared to the HFD group (Figure 3d-g). Specifically, the R+LP group showed a significant decrease in all biochemical markers compared to the LP and Q+LP groups while such differences were not observed in the groups administered *L. rhamnosus* ATCC 53103 (LR, Q+LR, and R+LR) (Figure 3d-g).

**Figure 1.**
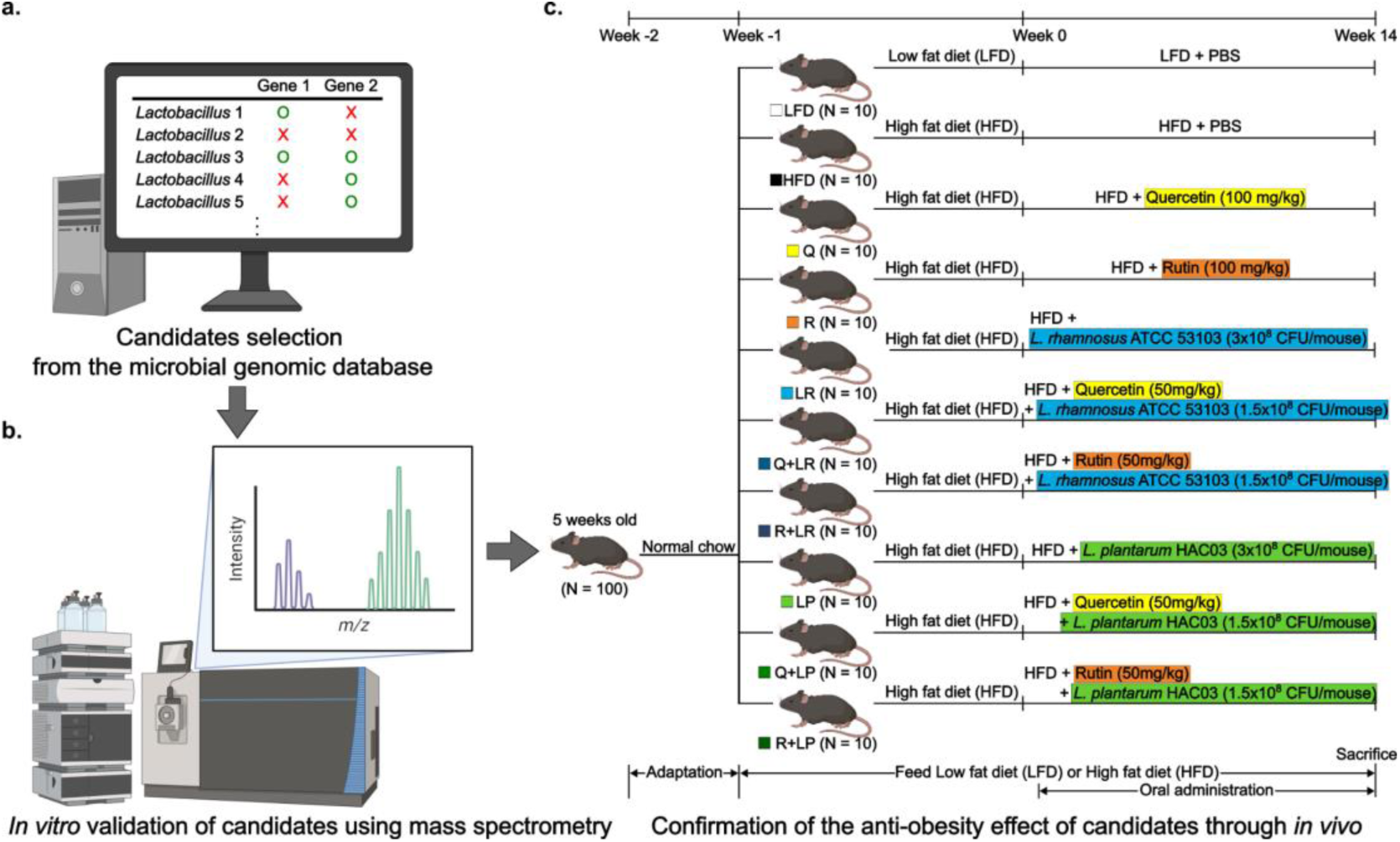
Experiment scheme. (a) Selecting candidates from the microbial genomic database. (b) Validating of candidates by analysing mass spectrometry results of the mixture of microbial candidates and flavonoids. (c) Confirming the anti-obesity effect of candidates through *in vivo* experiments. Some of the images included in this figure were created with assistance of BioRender.com.

**Figure 2.**
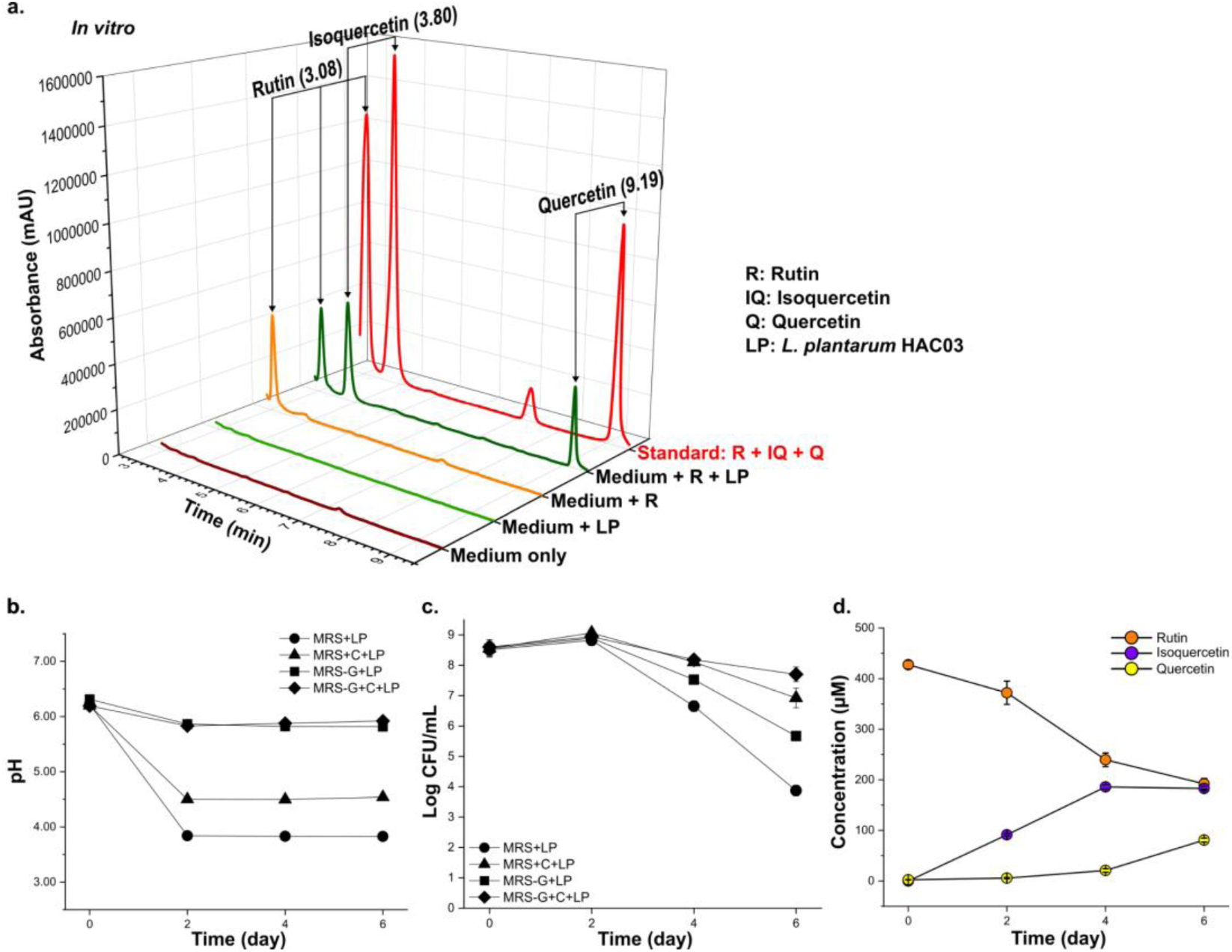
*L. plantarum* HAC03 hydrolyzes rutin into isoquercetin and quercetin. (a) HPLC-DAD-MS analysis of different media associated with *L. plantarum* HAC03 and rutin on day 6. (b-c) Among various growth media containing *L. plantarum* HAC03, the pH exhibited the smallest decrease, and the highest Colony-Forming Unit (CFU) count was observed in the glucose-free MRS broth with CaCO_3_ (MRS-g+C). (b) pH changes of *L. plantarum* HAC03 incubated on different media *in vitro*. (c) Growth curve of *L. plantarum* HAC03 incubated on different media *in vitro*. (d) The concentration changes of rutin, isoquercetin and quercetin in the *L. plantarum* HAC03 and rutin culture medium during the *in vitro*. The data are presented as means ± SD (n = 3).

**Figure 3.**
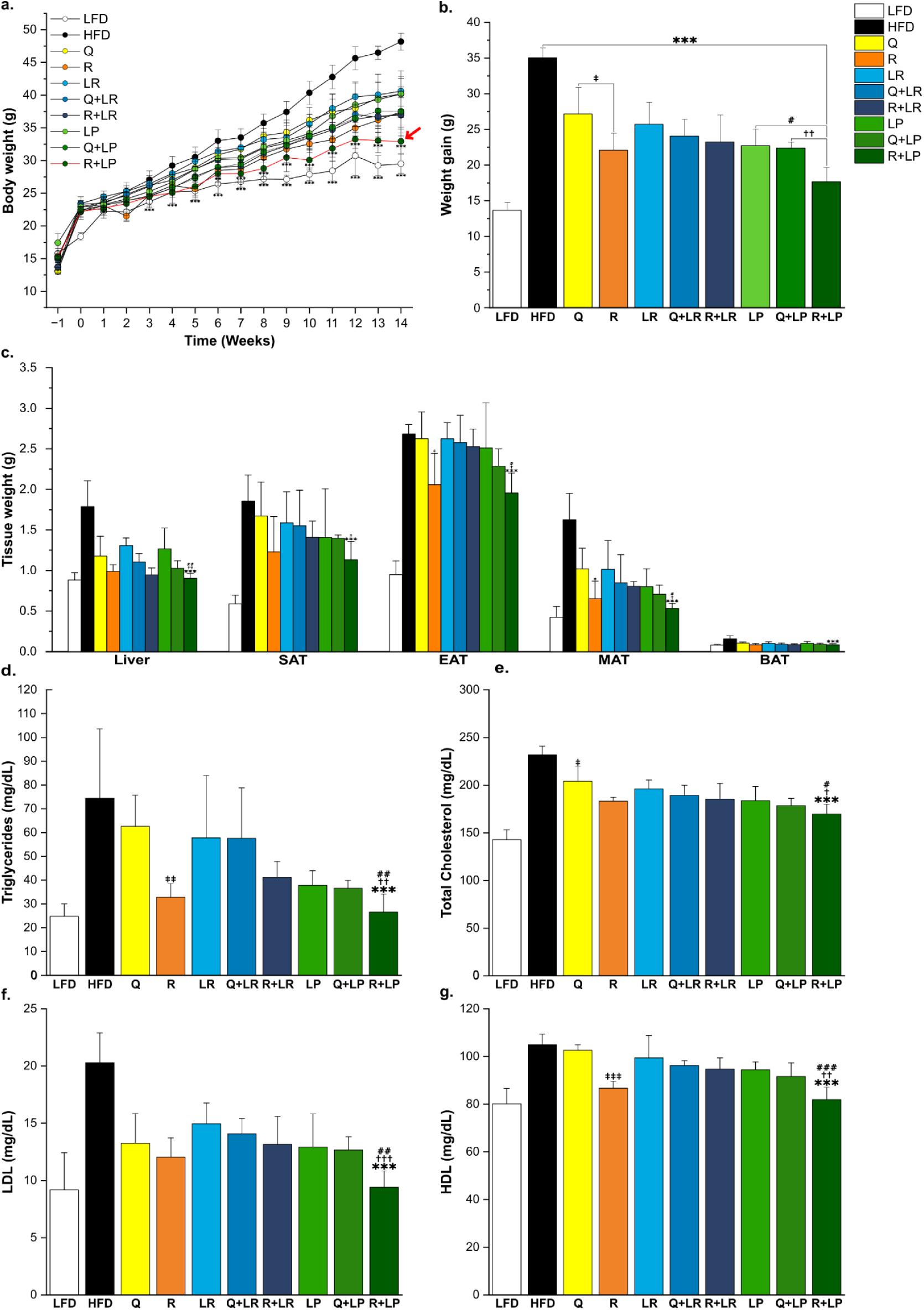
Combination of rutin and *L. plantarum* HAC03 has a synergistic effect on decreasing body weight. (a) Body weight changes during the animal experiment. (b) Weight gain during animal experiment (From Week −1 to Week 14). (c) Tissue weight after 14 weeks of treatment. Subcutaneous adipose tissue (SAT), epididymal Adipose Tissue (EAT), mesenteric Adipose Tissue (MAT), brown Adipose Tissue (BAT). (d–g) Biochemical indicators related to obesity in serum: (d) Triglyceride, (e) Total cholesterol, (f) LDL, (g) HDL. Low-fat diet; LFD, High-fat diet; HFD, Quercetin treatment; Q, Rutin treatment; R, *L. rhamnosus* ATCC 53103 treatment; LR, Quercetin and *L. rhamnosus* ATCC 53103 treatment; Q+LR, Rutin and *L. rhamnosus* ATCC 53103 treatment; R+LR, *L. plantarum* HAC03 treatment; LP, Quercetin and *L. plantarum* HAC03 treatment; Q+LP, Rutin and *L. plantarum* HAC03 treatment; R+LP. The data are presented as means ± SD (*n* = 10). One-way ANOVA with Tucky test was used for comparison with different groups. *** p < 0.001 between HFD and other groups. # p < 0.05, ## p < 0.01, ### p < 0.001 between LP group and R+LP group. † p < 0.05, †† p < 0.01, ††† p < 0.001 between Q+LP group and R+LP group. ‡ p < 0.05, ‡‡ p < 0.01, ‡‡‡ p < 0.001 between Q group and R group.

### Combination of *L. plantarum* HAC03 and rutin has a synergic effect on decreasing adipocyte size

Histological analyses were conducted on samples of four adipose tissues (subcutaneous adipose tissue; SAT, epididymal adipose tissue; EAT, mesenteric adipose tissue; MAT, brown adipose tissue; BAT) to systematically evaluate the anti-obesity effect. Compared to the quercetin treatment group (Q), the R group exhibited more decrease in adipocyte size in all adipose tissues (Figure 4b, d, f, and h). Furthermore, the R+LP group resulted in the greatest reduction in adipocyte count per unit area in all tissues compared with the HFD group (p-value < 0.001) (Figure 4b, d, f, and h). Particularly, in MAT among four adipose tissues, the adipocyte size of the R+LP group decreased by 73.05% compared to the HFD group, indicating the most substantial difference (Figure 4e, f). In contrast, the decrease was 24.78% in EAT (Figure 4c, d). The R+LP group exhibited a statistically significant decrease in three adipose tissues (SAT, MAT, and BAT) compared to those in LP or Q+LP groups. Among the groups associated with *L. rhamnosus* ATCC 53103 (LR, Q+LR, and R+LP), a significant reduction in adipocyte size was observed in all adipose tissues compared to that in the HFD group. However, no significant differences were observed between the groups in which the microbes were administerd alone or in combination with flavonoids.

**Figure 4.**
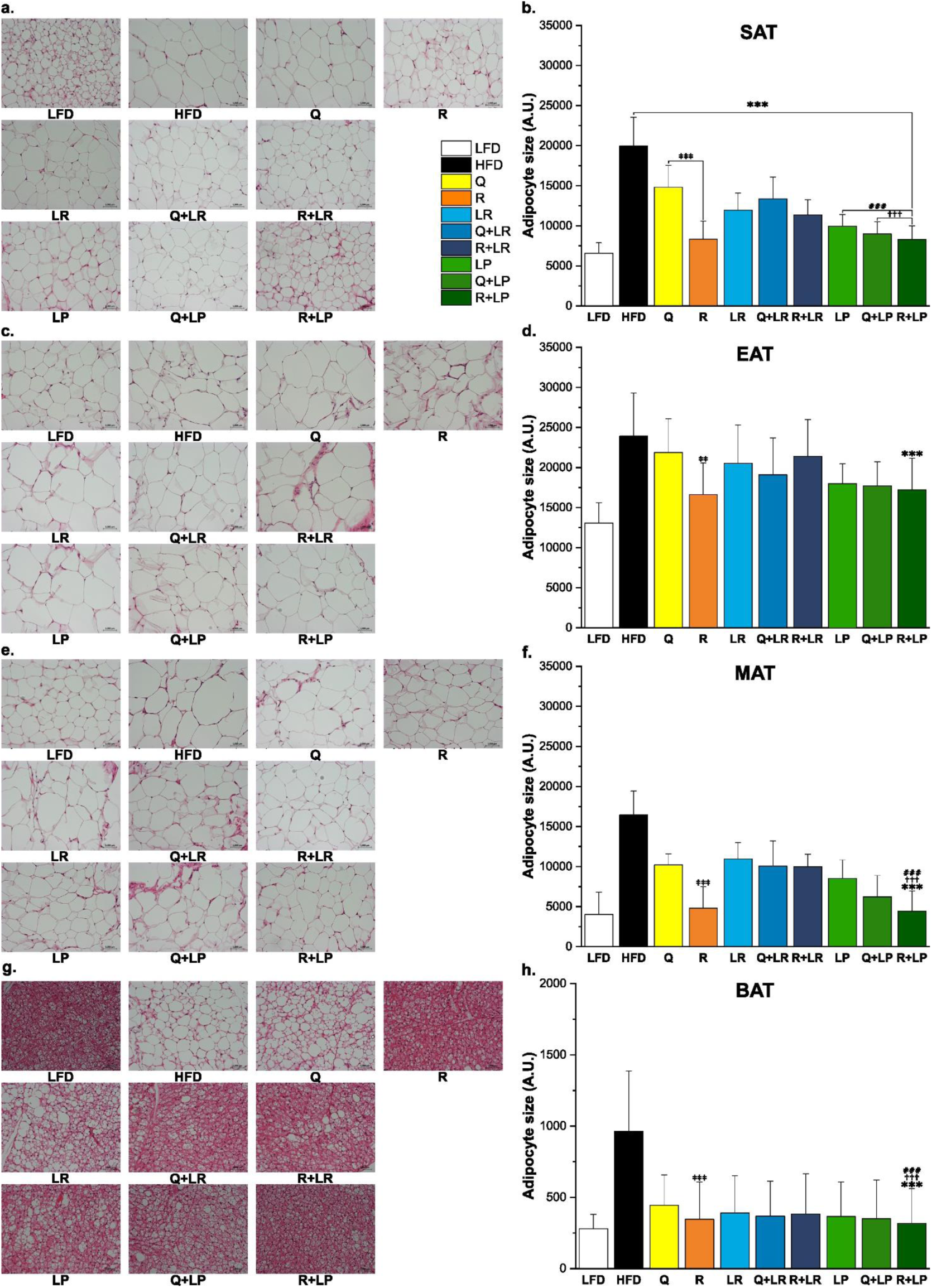
Combination of rutin and *L. plantarum* HAC03 has a synergistic effect on decreasing the adipocyte size. (a, c, e, g) Histological features of adipose tissue between the groups. Representative photomicrographs of adipose tissue sections stained with hematoxylin and eosin (×200) are shown. Subcutaneous adipose tissue (SAT), epididymal Adipose Tissue (EAT), mesenteric Adipose Tissue (MAT), brown Adipose Tissue (BAT). (b, d, f, h) Adipocyte size in adipose tissue between the groups. The data are presented as means ± SD (*n* = 5). One-way ANOVA with Tucky test was used for comparison with different groups. *** p < 0.001 between HFD and other groups. ### p < 0.001 between LP group and R+LP group. ††† p < 0.001 between Q+LP group and R+LP group. ‡‡ p < 0.01, ‡‡‡ p < 0.001 between Q group and R group.

### Combination of rutin and *L. plantarum* HAC03 induced anti-obesity effect by regulating gene expression related to obesity

The expression levels of obesity-related genes were examined in the four different adipose tissues. We measured the expression of genes related to fat synthesis (*FAS*, *ACC*, *PPARγ*, and *SREBP-1c*) and β-oxidation (*PPARɑ*, *PGC1ɑ*, *CPT1*, and *ACOX1*) in MAT, which is the adipose tissue in which the anti-obesity effect is the most pronounced. Compared to that in the HFD group, the expression of genes related to fat synthesis decreased in all groups, and the expression of genes related to β-oxidation increased (Figure 5a and b). The R group showed a decrease in the expression of fat synthesis-related genes *FAS* (p-value = 0.0013) and *PPARγ* (p-value = 0.0044) as well as an increased in the expression of the β-oxidation-related gene *CPT1* (p-value = 0.0033) compared to those in the Q group (Figure 5a). The R+LP group led to the most significant decrease in the expression of genes related to fat synthesis and the most significant increase in the expression of genes related to β-oxidation. The expression of all genes except for *ACOX1* was improved compared to LP (Figure 5b). Fat reduction involves a decrease in fat synthesis and increase in β-oxidation and plays a crucial role in thermogenesis in BAT and the browning of white adipose tissue.^30^ The R+LP group showed a significant increase in gene expression related to thermogenesis in BAT (*UCP1* and *PRDM16*), especially *UCP1* (p-value = 0.0001) compared with the HFD group (Figure 5c), and a significant increase in the expression of genes related to browning in SAT (*UCP1* and *PRDM16*) (Figure 5d).

**Figure 5.**
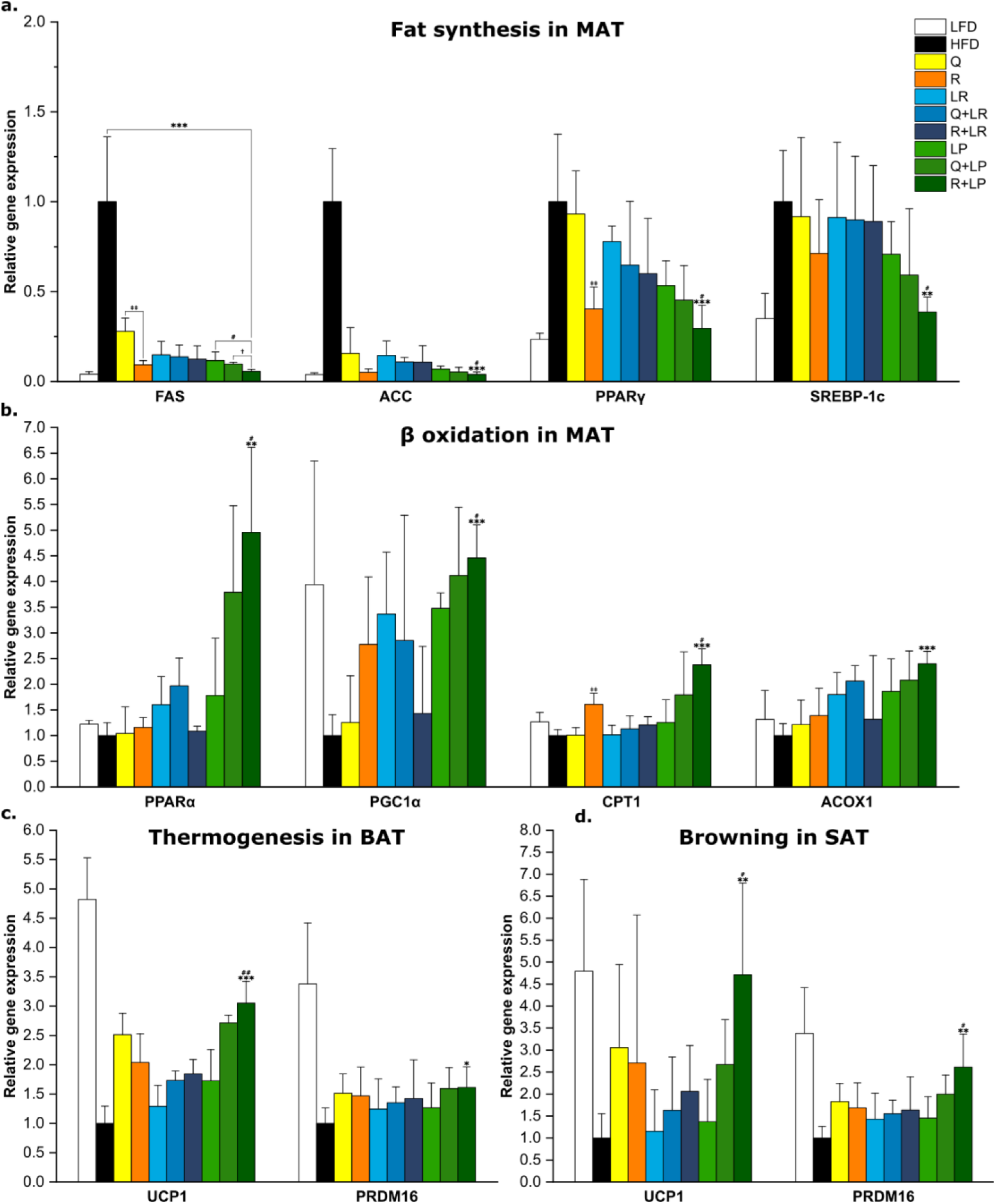
Combination of L. plantarum HAC03 and rutin induced an anti-obesity effect by regulating gene expression related to obesity. (a) Relative gene expression related to fatty acid synthesis in the MAT (*FAS*, *ACC*, *PPARɣ*, and *SREBP-1c*). (b) Relative gene expression related to β-oxidation in the MAT (*PPARɑ*, *PGC1ɑ*, *CPT1*, and *ACOX1*). (c) Relative gene expression related to thermogenesis in the BAT (*UCP1* and *PRDM16*). (d) Relative gene expression related to browning in the SAT (*UCP1* and *PRDM16*). The expression of all genes was normalized to that of β-actin. The data are presented as means ± SD (*n* = 8). One-way ANOVA with Tucky test was used for comparison with different groups. ** p < 0.01, *** p < 0.001 between HFD and other groups. # p < 0.05 between LP group and R+LP group. † p < 0.05 between Q+LP group and R+LP group. ‡‡ p < 0.01 between Q group and R group.

### Combination of rutin and *L. plantarum* HAC03 has a synergistic effect on alleviating insulin resistance

Obesity can induce insulin resistance, and this symptom can lead to various metabolic syndromes.^31^ We investigated whether the combination of rutin and *L. plantarum* HAC03 can alleviate insulin resistance, in addition to its impact on obesity. Oral glucose tolerance test (OGTT) at week 12 revealed that the R+LP group exhibited the second highest blood glucose levels after the HFD group at 20 min post-glucose administration (Figure 6d). However, as time progressed, the blood glucose levels rapidly decreased, reaching the second-lowest values after those in the LFD group at 60 min (Figure 6d). A comparison of the area under the curve (AUC) values confirmed that the R+LP group had the lowest value (Figure 6e). Furthermore, measurements of fasting blood glucose and insulin levels, and the calculated HOMA-IR index showed that the R+LP group had the lowest values after 14 weeks of oral administration (Figure 6a–c). Finally, the expression levels of *G6P* and *PEPCK*, which are related to gluconeogenesis in the liver, were the lowest in the R+LP group (Figure 6f, g). The combination of rutin and *L. rhamnosus* ATCC 53103 (R+LR) also showed a similar effect on insulin resistance.

**Figure 6.**
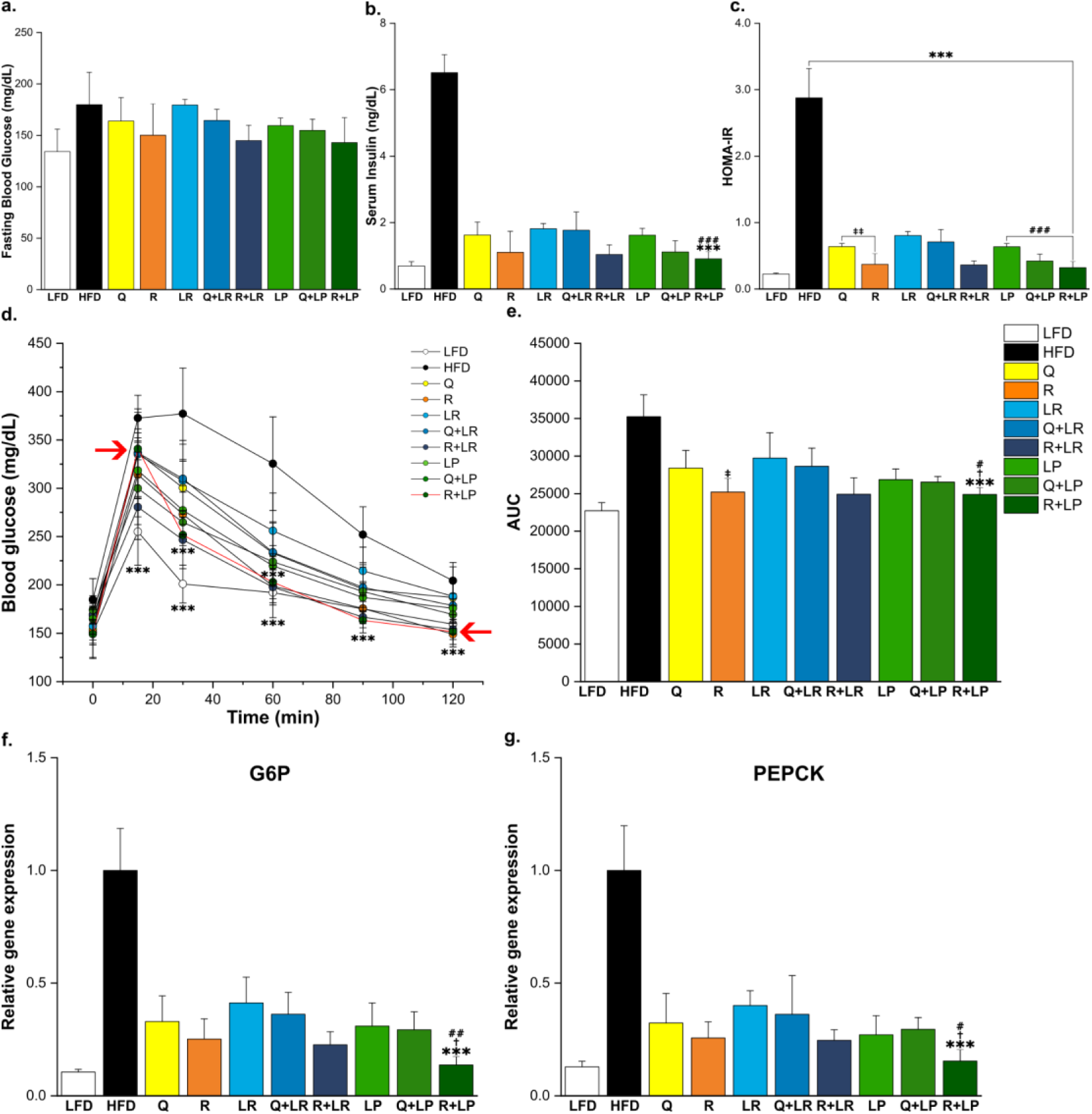
Combination of rutin and *L. plantarum* HAC03 has a synergistic effect on alleviating insulin resistance after 14 weeks of treatment. (a) Fasting blood glucose level (*n* = 10). (b) Fasting insulin level in serum (*n* = 10). (c) HOMA-IR index (fasting glucose (mg/dL) × fasting insulin (ng/dL) / 405) (n = 10). (d) Changes in blood glucose levels during the oral glucose tolerance test (n=10). (e) AUC curve. (f and g) Relative expression of gene related gluconeogenesis in liver (*G6P* and *PEPCK*). The expression of all genes was normalized to that of β-actin. The data are presented as means ± SD (*n* = 10). One-way ANOVA with Tucky test was used for comparison with different groups. *** p < 0.001 between HFD and other groups. # < 0.05, ## p < 0.01, ### p < 0.001 between LP group and R+LP group. † < 0.05, ††† p < 0.001 between Q+LP group and R+LP group. ‡ p < 0.05, ‡‡ p < 0.01 between Q group and R group.

### Combination of rutin and *L. plantarum* HAC03 has a synergistic effect on alleviating hepatic steatosis

Hepatic steatosis is a condition wherein a large amount of fat accumulates in the liver. It is the initial stage of nonalcoholic fatty liver disease (NAFLD) and it is closely related with obesity and diabetes.^32^ The previous results confirmed that the combination of rutin and *L. plantarum* HAC03 (R+LP) can significantly reduce liver weight compared to HFD group (Figure 3d). To investigate whether the combination of rutin and *L. plantarum* HAC03 is effective in alleviating hepatic steatosis, we observed the physiological features of liver tissues through histological analysis. The fat deposition in the liver significantly increased in the HFD group compared to that in the LFD group (Figure 7a). However, in all groups where the flavonoids or microorganisms were administered either individually or in combination, fat deposition significantly decreased. Particularly, the combination of rutin and *L. plantarum* HAC03 group (R+LP) exhibited the most significant reduction in fat deposition in the liver (Figure 7a). Furthermore, measurements of alanine aminotransferase (ALT) and aspartate aminotransferase (AST) levels in the serum, as indicators of liver damage, showed a significant decrease in the R+LP group compared to those in the HFD group (Figure 7b and c). In particular, both indicators exhibited significant reductions compared to the R and Q+LP groups. Finally, we determined the expression of genes related to fatty acid synthesis in the liver. The R+LP group showed the second-lowest expression levels after the LFD group (Figure 7d). Particularly, *FAS* and *SREBP-1c* exhibited significant reductions in the R+LP group, unlike in the *L. rhamnosus* ATCC 53103 treatment groups (Figure 7d).

**Figure 7.**
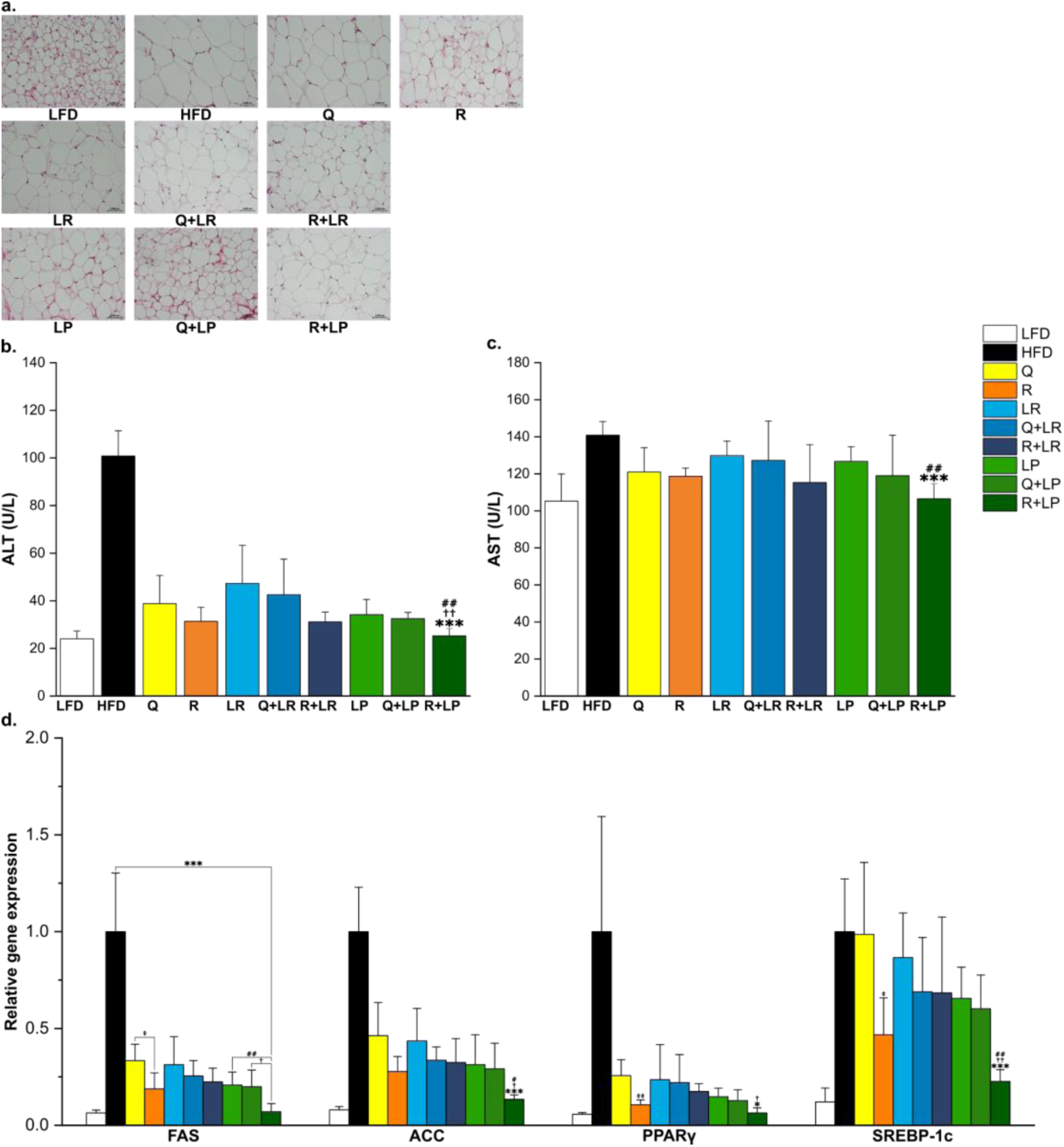
Combination of rutin and *L. plantarum* HAC03 has a synergistic effect on alleviating fatty liver. (a) Histological features of liver tissue between groups (n = 5). Representative photomicrographs of adipose tissue sections stained with hematoxylin and eosin (×200) are shown. (b and c) Biochemical indicators related to hepatocellular damage (n = 10): (b) alanine aminotransferase (ALT). (c) aspartate transaminase (AST). (d) Relative expression of fat synthesis gene in the liver (*FAS*, *ACC*, *PPARγ*, and *SREBP-1c*). The expression of all genes was normalized to that of β-actin. The data for quantitative PCR are presented as means ± SD (n = 10). One-way ANOVA with Tucky test was used for comparison with different groups. * p < 0.05, *** p < 0.001 between HFD and other groups. # p < 0.05, ## p < 0.01 between LP group and R+LP group. † p < 0.05, †† p < 0.01 between Q+LP group and R+LP group. ‡ p < 0.05, ‡‡ p < 0.01 between Q group and R group.

### Combination of rutin and *L. plantarum* HAC03 influenced the altered gut microbiota resulting from a high fat diet

To investigate the mechanism underlying the synergistic anti-obesity effect of rutin and *L. plantarum* HAC03, we first conducted a metagenomics analysis for faecal samples. The α-diversity showed a decrease in microbial richness and evenness in the HFD group compared to those in the LFD group. However, in the R+LP group, microbial richness and evenness, which was decreased in the HFD group, increased (Figure 8a–c). Furthermore, β-diversity differed between the groups. On the PCoA1 axis, which contained 25.32% of the total gut microbiota communities information, the R+LP group showed a recovery of gut microbiota to a level almost comparable to the low-fat diet group, despite being fed a high-fat diet. However, on the PCoA2 axis, encompassing 16.87% of the information, there was a substantial difference between the R+LP group and LFD group (Figure 8d). The relative distribution of *Lactobacillus* spp. increased in the LP and R+LP groups, in which *L. plantarum* HAC03 was administered (Figure 8f). Four bacteria in the R+LP group exhibited the most difference compared to the HFD group. *Eubacterium coprostanoligenes* abundance decreased compared to that in the HFD group, whereas *Lachnospiraceae* abundance increased. Additionally, *Oscillospiraceae* and *Eggerthellaceae* abundance increased in the R+LP group compared to that in the HFD group (Figure 8e-g). The changes in the distribution of the four gut bacteria under the given conditions are consistent with previously reported findings.^33–37^

**Figure 8.**
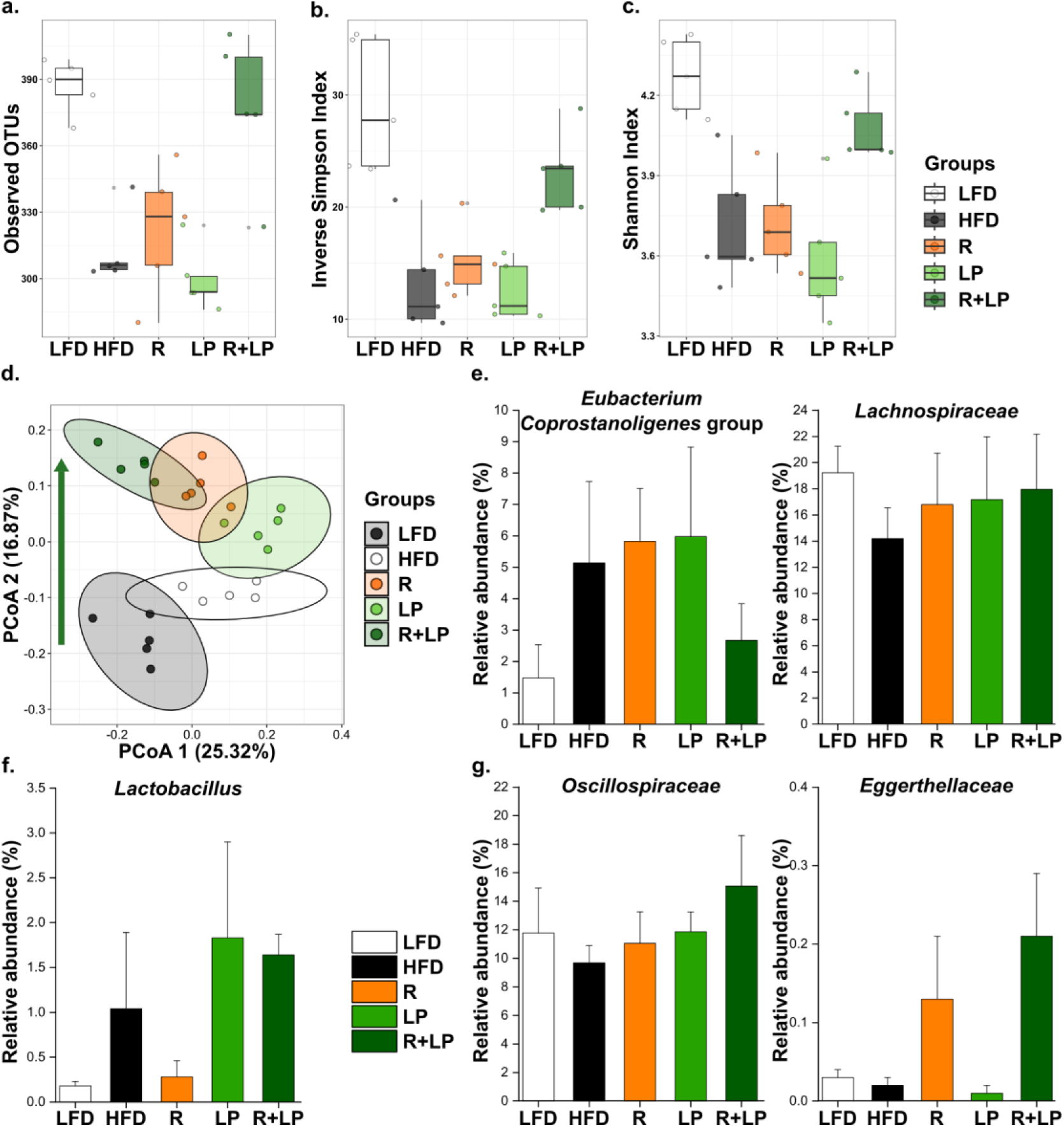
The combination of rutin and *L. plantarum* HAC03 affect gut microbiota modulation. ɑ-diversity: (a) Observed operational taxonomic units (OTUs), (b) Inverse simpson index, (c) Shannon index. (d) β-diversity. (e) Relative distribution of specific gut microbiota groups associated with obesity. (f) Relative distribution of *Lactobacillus* spp. (g) Relative distribution of specific gut microbiota groups associated with flavonoids metabolism. All gut microbiota analyses were performed using fecal samples. The data are presented as means ± SD (n = 5).

### Combination of rutin and *L. plantarum* HAC03 has a synergic effect on the localization of *L. plantarum* in the ileum

We investigated the relative abundance of *L. plantarum* among the five experimental groups in the three intestinal regions (duodenum, ileum, and colon) using quantitative PCR. To compare the amount of *L. plantarum* across the different testing groups in the three intestinal regions, we normalized the quantity of *L. plantarum* in the HFD group to 1 for the three intestinal regions (Figure 9). In the colon, all testing groups did not exhibit significant increases compared to the HFD group (Figure 9d). However, in the ileum, a substantial increase was observed when rutin and *L. plantarum* HAC03 were co-administered while the LP alone group did not show any significant change compared to the HFD group (Figure 9c). We also included faecal samples in this analysis. The qPCR results, measuring the relative abundance of *L. plantarum* in each group, matched the metagenomic sequencing results obtained earlier (Figure 8f and 9e). There are marginal increases in both LP and R+LP groups compared to the HFD group, but the significant difference between the LP and the R+LP group confirmed in the ileum region was not observed (Figure 9e).

**Figure 9.**
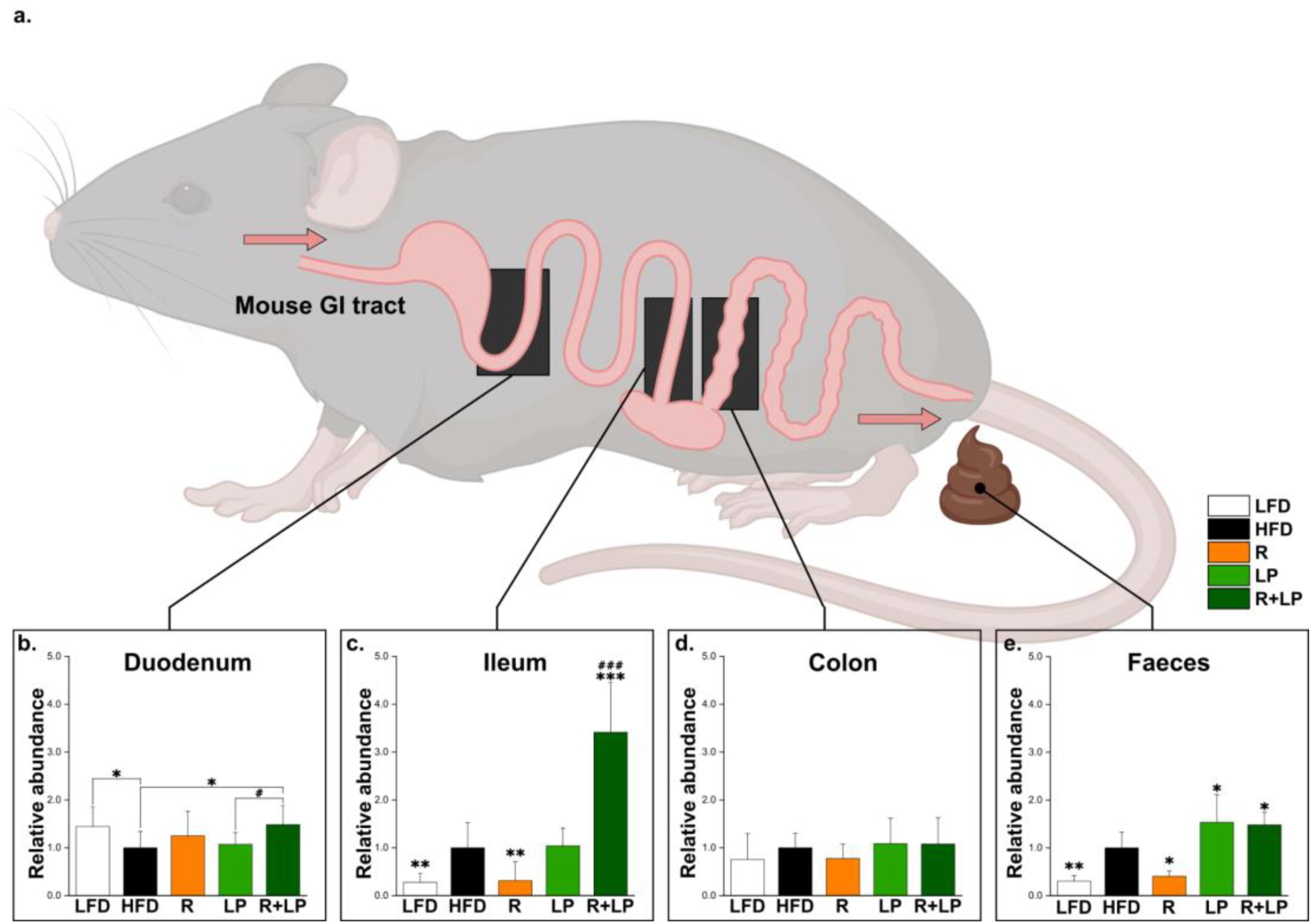
The combination of rutin and *L. plantarum* HAC03 affect the *L. plantarum* localization in the gut. (a) Location within the intestine where the samples were collected for the experiment (duodenum, ileum, colon, and faeces). (b–e) Relative composition of *L. plantarum* in different locations in the gut: (b) duodenum, (c) ileum, (d) colon, and (e) faeces. The relative composition of *L. plantarum* was normalized with the 16S rRNA gene of universal bacteria. The data for quantitative PCR are presented as the means ± SD (n = 5). One-way ANOVA with Tucky test was used for comparison with different groups. * p < 0.05, *** p < 0.001 between HFD and other groups. # p < 0.05, ### p < 0.001 between LP group and R+LP group. The mouse and mouse GI tract images in this figure were created using BioRender.com.

## Discussion

Flavonoids are a subset of phytochemicals produced by plants such as vegetables and fruits. Rutin is one of the most abundant flavonoids and has been extensively studied.^38,39^ Numerous clinical trials have confirmed that oral administration of plant-based flavonoids is extremely safe with no side effects.^40^ Moreover, flavonoids exert an anti-obesity effect by controlling fat synthesis and adipocyte lifespan, inducing thermogenesis to increase energy expenditure in the body, regulating nutrient absorption, reducing food intake and modulating the gut microbiota.^8,42,43^ Specifically, through *in vitro* experiments, rutin has been confirmed to exhibit dose-dependent increases in gene expression associated with anti-obesity and anti-diabetes effect at the cellular level.^44^ Additionally, *in vivo* experiments utilizing a DIO mouse model demonstrated that oral administration of rutin for seven weeks resulted in a decrease in the weight gain rate, providing further evidence for the anti-obesity effect of rutin.^45^ However, despite being one of the most abundant flavonoids in fruits and vegetables, rutin’s low bioavailability hinders its optimal utilization as a health supplement.^46^

We hypothesized that increasing rutin’s bioavailability could enhance its anti-obesity properties. To validate this hypothesis, our focus turned to *Lactobacillus* species, a group of microorganisms that show the capacity to hydrolyze rutin in the human and murine intestines.^47–49^ In addition, to confirm whether the increase in the anti-obesity effect that can be caused by the combination of the two is additive or synergistic, the total amount of all substances administered to the test groups was kept the same. If combining rutin and a *Lactobacillus* strain results in effects similar to those observed when each is administered alone, it indicates an additive effect (i.e. 0.5 + 0.5 = 1). However, if more significant changes occur despite halving the concentrations of each, it suggests a synergistic interaction between them (i.e. 0.5 + 0.5 > 1).

We found that despite no changes in the average feed intake among groups, *L. plantarum* HAC03 synergistically increases the anti-obesity effect when co-administered with the flavonoid rutin and the *in vitro* and *in vivo* experiments suggested the following conclusions.

First, although rutin’s bioavailability is generally lower than that of quercetin in the human and murine intestine ^9,50^, the rutin treatment group (R) exhibited a significant reduction in body weight gain compared to that of the quercetin treatment group (Q) (Figure 3b). In further analysis, the R group showed reduced adipocyte size, downregulated expression of genes contributing to weight gain, and increased expression of genes associated with energy expenditure in comparison to those of the Q group (Figures 4 and 5). In the early stages of our study, we anticipated that the Q group would exhibit a superior anti-obesity effect compared to the R group. However, the experimental results were contrary to our expectations. We inferred that the gut microbiota of the mice used in this study play a role in generating these differences in anti-obesity effect. Gut microbiota are essential for the absorption of rutin into the body.^51^ Gut microbiota can sequentially convert rutin into isoquercetin, quercetin, phenolic acids.^52^ The fact that the group treated with rutin alone (R group) showed similar or even superior anti-obesity effect compared to the group treated with pure quercetin (Q) suggests the possibility that other rutin derivative(s) produced by gut microbiota have more potent anti-obesity effect than quercetin. Additionally, gut microorganism(s) responsible for producing such potent rutin derivative(s) may be the third species rather than mouse-derived *L. plantarum*. This is suggested by the significant decrease of the mouse-derived *L. plantarum* in the ileums of the rutin-only treated group (R) compared to that in the other groups (Figure 9c). Note that the mouse-derived *L. plantarum* is different from the plant-derived *L. plantarum* HAC03 investigated in this study.

The combination of rutin and *L. plantarum* HAC03 (R+LP) that exhibited a synergistic anti-obesity effect induces numerous changes in gut microbiota, gene expression in the adipose tissues, and blood serum composition. Metagenomic analysis showed that certain gut bacteria, namely *Eubacterium coprostanoligenes*, *Lachnospiraceae, Oscillospiraceae,* and *Eggerthellaceae,* exhibited changes in response to the combination treatment in the R+LP group. For example, the abundance of *Eubacterium coprostanoligenes* significantly decreased compared to that in the HFD group, reaching levels comparable to those observed in the LFD group (Figure 8e). This result is in line with the increase of *Eubacterium coprostanoligenes* observed in humans with obesity and type 2 diabetes.^33^ While statistical significance wasn’t achieved, there was a tendency for *Lachnospiraceae* to increase in the R+LP group, supporting previous report that *Lachnospiraceae* are negatively associated with overweight/obese individuals.^34^ In addition, the increase in *Oscillospiraceae* and *Eggerthellaceae* compared to the HFD group and R groups (Figure 8g) is also consistent with the elevated levels of these bacteria observed in humans consuming flavonoid-zor polyphenol-rich foods.^35–37^ The high consistency between these findings and prior human research indicates that the mouse-based experiments in this study can be valuable models for human obesity research. While it remains unclear whether these changes in gut microbiota are the cause or result of the strong anti-obesity effect observed in the R+LP group, the correlation between the two is evident.

We found that rutin plays an important role in facilitating the colonization of *L. plantarum* HAC03 in the ileum. To explore the intraintestinal localization of *L. plantarum* in the testing groups, we conducted quantitative PCR targeting the conserved genomic region specific to *L. plantarum* species. In our experiment, the LP group (the group that added only *L. plantarum* HAC03 to high-fat feed) did not exhibit any increase of *L. plantarum* in ileum compared to that in the HFD group. The high-fat diet itself does not appear to be helpful for the colonization of the fermented Kimchi-derived *L. plantarum* HAC03 in the mouse intestine. However, when rutin was co-administered with *L. plantarum* HAC03, the amount of *L. plantarum* was significantly increased in the ileum compared to other groups (Figure 9c). Interestingly, administering rutin alone without *L. plantarum* HAC03 resulted in a substantial decrease in the pre-existing *L. plantarum* in the mouse ileum. This suggests that rutin could function as an effective prebiotic for the settlement of the fermented plant-derived *L. plantarum* HAC03 *in vivo*, while potentially acting as an antimicrobial agent for the indigenous *L. plantarum* derived from mice raised on high-fat feed.

The significant changes resulting from the co-administration of rutin and *L. plantarum* HAC03 (R+LP) were also observed in the gene expression within three different adipose tissues (MAT, BAT and SAT). Among the 10 obesity-related marker genes monitored in this study, the expression levels of 5 genes in different tissues showed synergistic increases or decreases exclusively in the R+LP group compared to them in the HFD, R and LP groups (Figure 5). These 5 genes are involved in the following biochemical reactions: fatty acid synthesis (FAS), β-oxidation (PPARα, PGC1α, CPT1), and thermogenesis in borwn adipose tissue (UCP1). On the other hand, no synergistic anti-obesity effect was observed when treated with rutin and the *L. rhamnosus* strain lacking the ability to decompose rutin. This result suggests that the rutin derivative(s) produced by the R+LP combination inhibit or accelerate the biochemical reactions mentioned above, contributing to the synergistic anti-obesity effect.

So, what substance(s) and mechanism(s) of action could induce this synergistic anti-obesity effect? Based on our experimental results, we propose a model (Figure 10). In this model, isoquercetin, produced in the ileum by *L. plantarum* HAC03 from orally administered rutin, is a leading compound responsible for achieving synergistic anti-obesity effect in the R+LP group. This rutin-derivative, known for its high bioavailability and superior anti-obesity effect, is predominantly absorbed in the small intestine by enzymes and transporters.^16,23,45^ Our *in vitro* experiment showed that *L. plantarum* HAC03 rapidly converted rutin into isoquercetin from the early stages of the experiment. On the other hand, the conversion to quercetin occurred very slowly (Figure 2d). If *L. plantarum* HAC03 follows the similar pattern *in vivo*, the *L. plantarum* strain settled in the ileum will quickly convert orally administered rutin into isoquercetin, and the newly produced isoquercetin will be effectively absorbed in the ileum before further converting to quercetin. The fact that no synergy or a slight synergy in various biomarkers was observed in the Q+LP group compared to R+LP suggests that isoquercetin, the precursor to quercetin in the rutin biotransformation, will play a key role in enhancing the anti-obesity effect. There is a possibility that isoquercetin absorbed in the small intestine is distributed directly to various tissues through the bloodstream, or is converted to substances other than quercetin by enterocytes (major intestinal cells) or other gut microorganisms, thereby producing enhanced anti-obesity effect.

**Figure 10.**
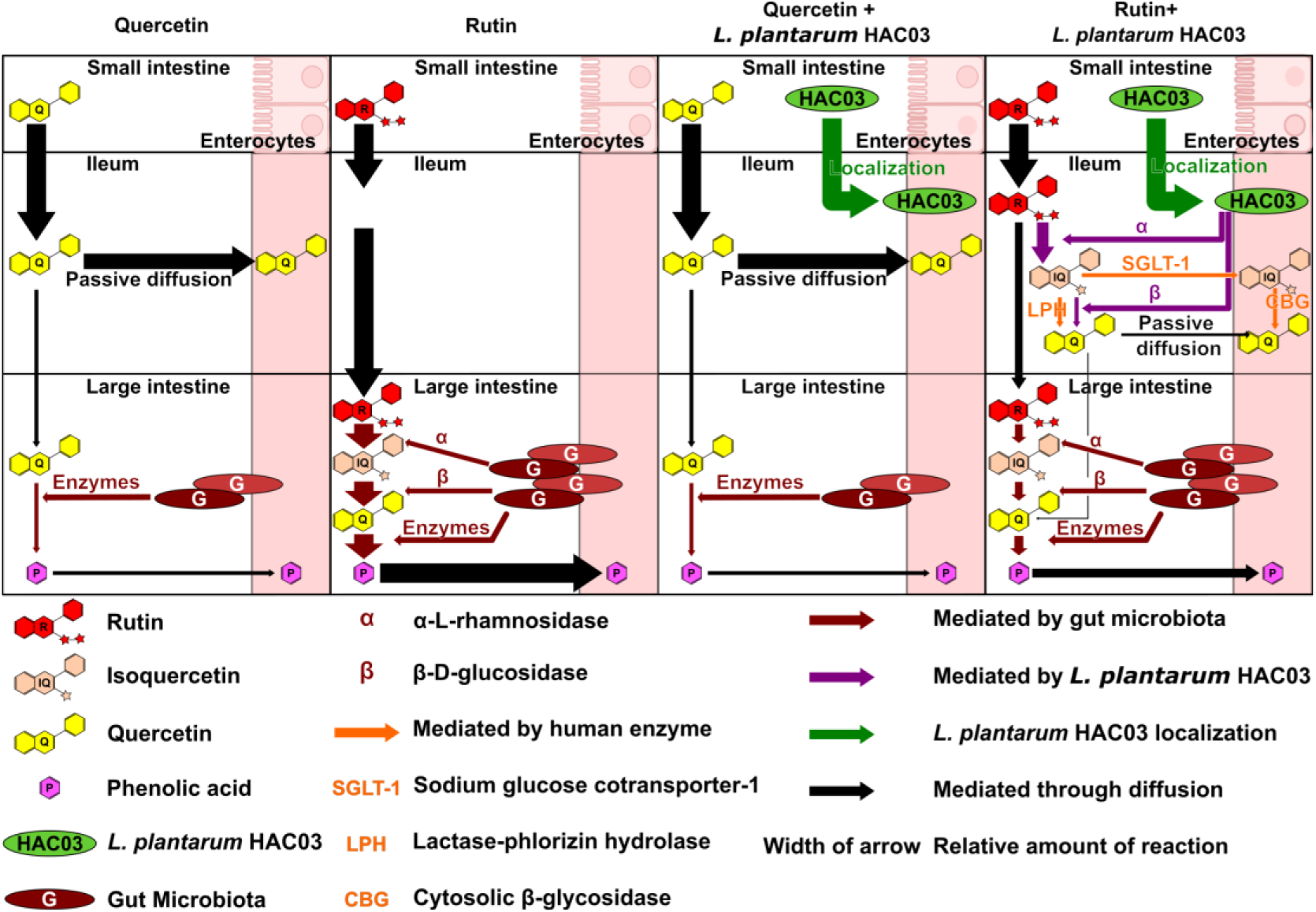
A model of enhanced anti-obesity effect when *L. plantarum* HAC03 and rutin were treated in combination. Intestinal metabolism following the oral administration of each substance is proposed. Quercetin is primarily absorbed in the small intestine through passive diffusion, whereas rutin undergoes hydrolysis in the large intestine before absorption, mainly as phenolic acid. Co-administration of quercetin with *L. plantarum* HAC03, localized in the ileum has no significant impact on quercetin absorption. Conversely, co-administration of *L. plantarum* HAC03 with rutin enhances the rutin bioavailability in the ileum by hydrolyzing it into isoquercetin and quercetin. This combination modifies the gut microbiota in the large intestine, leading to increased flavonoid metabolism and an enhanced anti-obesity effect.

In this study, we discovered a novel combination that showed a synergistic anti-obesity effect, systematically analyzed the *in vivo* effect of this combination. We proposed the mechanism underlying the synergistic anti-obesity effect. These results showed the possibility of this combination as a safe and highly effective anti-obesity solution. Furthermore, the effective reduction of mesenteric adipose tissue (Figure 4f), which belongs to visceral fat highlights the potential of this combination as a health supplement inhibiting the accumulation of visceral fat, which can cause inflammation and various chronic diseases.^53^

Finally, we want to emphasize the importance of spatial analysis of the gut microbiome. In this study, we divided the intestine into three regions and observed the distributions of *L. plantarum* among the experimental groups within these regions. The results showed significant differences in the distribution between the regions (Figure 9b-d). However, the distributions observed in faecal samples (Figure e), commonly used for metagenomic analysis, proved inadequate in capturing the dynamic variations dependent on the intestinal location, especially in the small intestine. This tendency was particularly pronounced in the R+LP group. We confirmed a remarkable increase in *L. plantarum* in the ileum of mice that were orally administered both rutin and *L. plantarum* HAC03. However, such significant differences were not observed in faecal samples. Several studies have also indicated that while faecal samples may exhibit a distribution similar to that of the colonic bacterial community, they diverge from the bacterial community in the small intestine.^54–56^ If the regional analysis of the intestine had not been conducted in this study, we would not have discovered that rutin serves as an effective prebiotic for the colonization of *L. plantarum* HAC03 in the ileum. Additionally, we would not have been able to propose the model suggesting that orally administered rutin is converted into isoquercetin by the *L. plantarum* HAC03 settled in the ileum and is effectively absorbed in the ileum, emphasizing the significance of spatial analysis in the gut for uncovering more detailed mechanisms of intriguing phenomena, including the synergistic anti-obesity effect.

## Materials and methods

### Bacterial Strains, Culture Conditions and Natural Product

*L. plantarum* HAC03 strain, originated from fermented white Kimchi in South Korea^57^, was provided by the Handong Global University Office of Industry-Academic Research (Pohang, Gyeongsangbuk-do, Republic of Korea). *L. rhamnosus* ATCC 53103 strain served as the positive control for anti-obesity effect and was purchased from American Type Culture Collection (ATCC) (Manassa, VA, USA). Both strains were grown in MRS broth (Difco Laboratories INC., Franklin Lakes, NJ, USA) at 37 ℃. For administration to the mice, the bacterial cells were dissolved in 1× Phosphate-buffered saline (PBS) at a concentration of 3 × 10^9^ CFU per mouse. Quercetin dihydrate and rutin trihydrate were obtained from DAEJUNG Chemicals & Metals (Siheung, Gyeonggi-do, Republic of Korea).

### Selection of *Lactobacillus* candidates

Microbial candidates were selected using the KEGG ORTHOLOGY (version 101.0). Within the KEGG ORTHOLOGY database, bacteria belong to the genera *Lactobacillus*, *Lacticaseibacillus*, and *Lactiplantibacillus* were identified based on the presence of at least one of the following genes: *RamA* (K05989), *bglX* (K05349), and *bglA* (K01223). Further filtering was used to select the bacteria that possessed both *RamA* (K05989) and either *bglX* (K05349) or *bglA* (K01223).

### Rutin biotransformation by *Lactobacillus* strains

A bacterial culture with a concentration of 1 × 10^10^ CFU was inoculated into a medium with 20 mL glucose free MRS and with 0.5% CaCO_3_ (Biosesang, Yongin, Gyeonggi-do, Republic of Korea) containing 0.5 mM rutin to investigate the capability of *L. plantarum* HAC03 and *L. rhamnosus* ATCC 53103 to convert rutin into their aglycone forms (isoquercetin and quercetin). The cultures were incubated at 37 ℃ for six days. The pH value and growth curves of the cultures were monitored. After incubation, 10 mL of dimethyl sulfoxide (DMSO) (Biosesang, Yongin, Gyeonggi-do, Republic of Korea) were added to the samples. The samples were filtered using a 0.2 µm syringe filter (Sartorius, Göttingen, Land Niedersachsen, Germany), after which the filter was rinsed with an additional 5 mL of DMSO. The sample was freeze-dried by using a FD8508 (IlshinBioBase Co. Ltd., Dongducheon, Gyeonggi-do, Republic of Korea). The dried samples were reconstituted in an appropriate volume of DMSO to prepare the samples for HPLC-DAD-MS measurements.

For quantitative analysis using HPLC-DAD-MS, the reference compounds (quercetin, isoquercetin and rutin) were dissolved in DMSO to form a 1 mM stock solution and used at 1 μM to 1 mM concentrations for HPLC analysis. The fingerprint was determined by HPLC-DAD analysis and validated by mass spectrometry. The concentrations of compounds were analyzed by DIONEX ultimate 3000 HPLC system, consisting of a pump (HPG-3200RS), sampler (WPS-3000), column oven (TCC-3000), and diode array detector (VWD-3100) equipped with Thermo LTQ XL mass spectrometer. A gradient elution was performed in a Thermo Scientific Acclaim 120 C18 column (120 Å, 150 mm × 2.1 mm, 5 μm pore size). The gradient system consisted of mobile phase A (0.1% (v/v) acetic acid) and mobile phase B (acetonitrile). All solutions were degassed and filtered through a 0.45 μm pore size filter prior to analysis. The initial conditions were as follows: 25% of eluent B with a linear gradient to 35% from 0 to 5 min, followed by a linear gradient to 100% of eluent B at 10 min, which was maintained for 2 min. A proportion of the mobile phase was returned to the initial concentration with a linear gradient at 15 min and allowed to equilibrate pressure for another 5 min before the injection of another sample. The total runtime of each sample was 20 min. The flow rate was 0.2 mL/min and the sample injection volume was 10 μL. The HPLC system was maintained at 25 ℃ for the analysis and the UV detector was set at 264 nm. Under these chromatographic conditions, it was possible to detect distinct retention time (RT) of rutin, isoquercetin and quercetin without overlap and they were confirmed by monitoring the deprotonated molecular ions [M-H]-at m/z 609, 463 and 301 respectively in the mass spectrum.

### Animal experiments

Animal experiments were conducted in compliance with protocols approved by the Committee on the Ethics of Animal Experiments at Handong Global University (permit number: 20221026-13). Five-week-old C57BL/6 male mice obtained from Saeronbio Inc (Uiwang, Gyeonggi-do, Republic of Korea) were housed at 23 ± 1 ℃, 60 ± 5% humidity, and a 12 hour light/dark cycle. After one week of adaptation, the mice were divided into 10 groups (*N* = 10 per group). Table 1 provides a detailed description of the groups. One week prior to the oral administration of bacteria or flavonoids, either LFD (10% kcal from fat, D12450J, Research Diets Inc., New Brunswick, NJ, USA) or HFD (60% kcal from fat, D12492, Research Diets Inc., New Brunswick, NJ, USA) was administered to the mice. Subsequently, the respective LFD or HFD diets were provided in conjunction with either bacteria (*L. rhamnosus* ATCC 53103 or *L. plantarum* HAC03) at a dose of 3×10^8^ CFU/mouse or flavonoids (Quercetin or Rutin) at a dose of 100 mg/kg over 14 weeks. The administrations were conducted individually or in combination based on the designated experimental groups. On the last day of the experiment, the mice were euthanized using CO_2_ gas. The organs (liver, small intestine, colon, subcutaneous adipose tissue; SAT, epididymal adipose tissue; EAT, mesenteric adipose tissue; MAT, and brown adipose tissue; BAT) were collected and stored at −80 ℃ until further analyses.

**Table 1.**
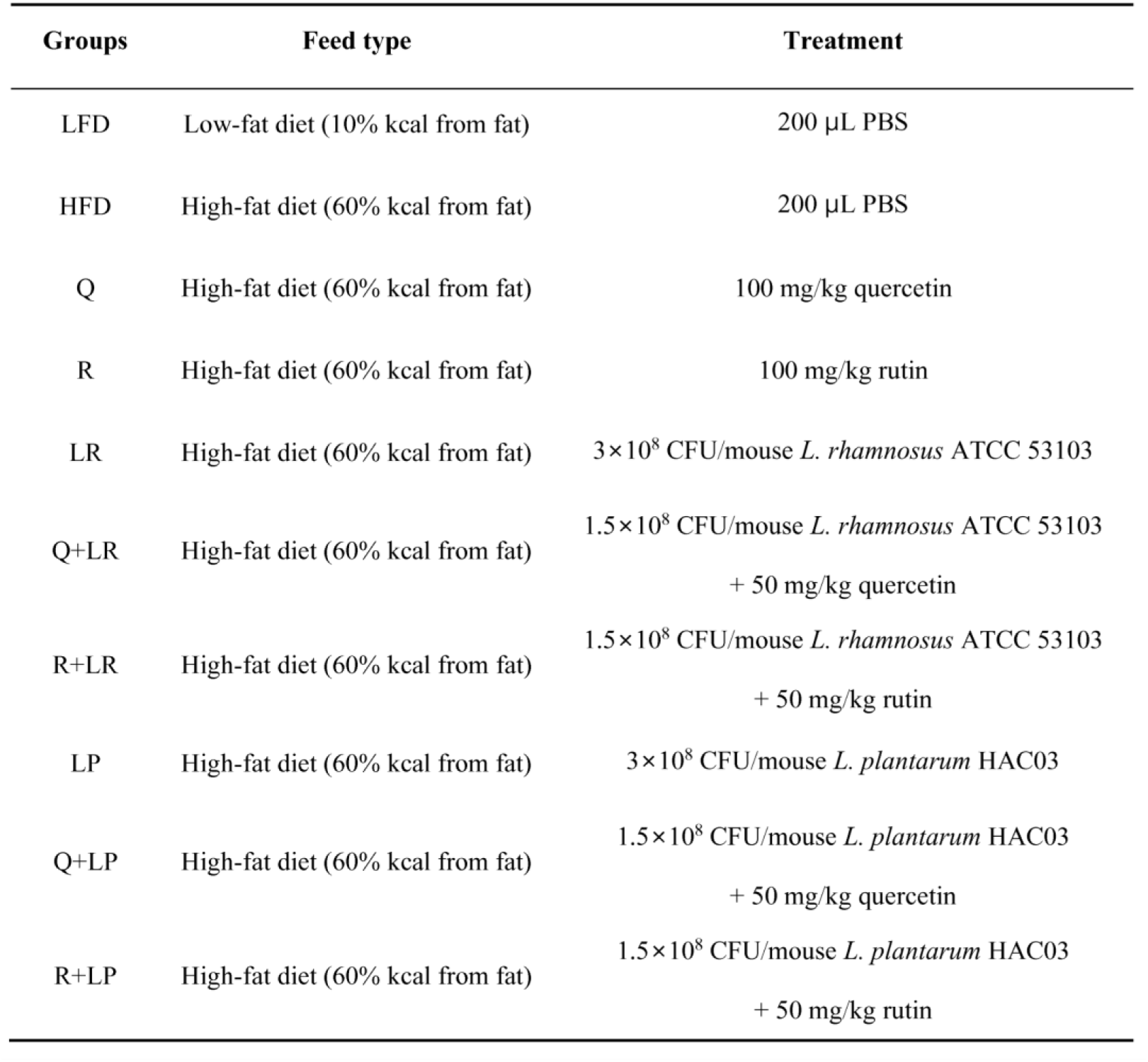
Animal experimental groups.

### Oral glucose tolerance test

After 12 weeks of treatment, the mice fasted for 6 hours with unrestricted access to water. Following this fasting period, glucose (Biosesang, Yongin, Gyeonggi-do, Republic of Korea) was orally injected at concentration of 2g/kg body weight. Blood samples were collected from tail bleeds at baseline and 15, 30, 60, 90, and 120 min after glucose injection, and glucose levels were measured using GlucoDr auto AGM-4000 (Allmedicus, Anyang, Gyeonggi-do, Republic of Korea).

### Serum analysis

Blood was obtained via cardiac puncture and collected in a blood collection tube with lithium heparin (Becton, Dickinson and Company, Franklin Lakes, NJ, USA). The collected blood was then centrifuged for 15 minutes at 2000 × g, and 4 ℃ to separate the serum. Biochemical parameters in the serum, including total cholesterol, triglycerides, alanine aminotransferase (AST), aspartate aminotransferase (ALT), low density lipoprotein (LDL), and high density lipoprotein (HDL) were analyzed using a 7180 Clinical Analyzer (Hitachi High-Tech, Tokyo, Japan).

### Histological analysis

Part of the adipose tissue was fixed in 10% v/v formalin/PBS solution for histological analysis. It was embedded in paraffin for staining with hematoxylin and eosin. Images were captured using a ZEISS Axio Imager 2 (Carl Zeiss Co. Ltd., Land Baden-Württemberg, Germany) at a magnification of 200×. The surface area of adipocytes were quantified using ImageJ software with an Adiposoft plug-in, following the developer’s instructions.

### Real-time polymerase chain reaction (RT-PCR)

The total RNA was extracted from liver and adipose tissues using the TRIzol Reagent (Invitrogen, WALTHAM, MA, USA). cDNA was synthesized using a SuperiorScript III cDNA Synthesis kit (Enzynomics Inc., Daejeon, Republic of Korea). Quantitative PCR (qPCR) was carried out on an Applied Biosystem StepOnePlusTM Real-Time PCR system (Applied Biosystems, WALTHAM, MA, USA) using a TOPrealTM qPCR 2X PreMIX (SYBR Green with high ROX) kit (Enzynomics Inc., Daejeon, Republic of Korea). All primers used for the qPCR were synthesized by Macrogen (Seoul, Republic of Korea). The sequences of all qPCR primers used in this study are listed in Supplementary Table 2. The results are presented as means ± S.D. normalized to β-actin expression using the ΔΔCt method, in which the HFD group was used as the reference group.

### Gut microbiota analysis

A Metagenomic analysis was carried out using the following protocol. First, DNA extraction from faecal samples was performed with the AccuStool DNA Preparation kit (AccuGene, Incheon, Republic of Korea) following the manufacturer’s instructions. The hypervariable V3-V4 region of the 16S-rRNA gene was amplified from the DNA extracts through 25 PCR cycles using the KAPA HiFi HotStart ReadyMix (Roche, Basel, Swiss) and barcoded fusion primers 341f/805r containing Nextera adaptors.^58^ The PCR products (∼428 bp) were purified with HiAccuBeads (AccuGene, Incheon, Republic of Korea). Amplicon libraries were pooled at equimolar ratio, and the pooled libraries were sequenced on an Illumina MiSeq system using the MiSeq Reagent Kit v3 with 600 cycles (Illumina, San Diego, CA, USA). All raw datasets were denoised by correcting the amplicon errors and used to infer the exact amplicon sequence variants (ASVs) using DADA2 v1.16.^59^ The SILVA release 138 rRNA reference database was used to create a Naïve Bayes classifier to classify the ASVs obtained from DADA2.^60^ Downstream analyses of quality- and chimera-filtered reads were performed using the QIIME2-2022.8 software package.^61^ Each sequences obtained from the DADA2 datasets was assigned to a taxonomy with a threshold of 99% pairwise identity using QIIME2 workflow scripts and the SILVA release 138 rRNA reference database classifier. Gut microbiota analysis, encompassing alpha diversity, beta diversity, and relative abundance of the gut microbiota was performed using Microbiome.

### Spatial mapping of Lactiplantibacillus plantarum

The localization of orally administered *L. plantarum* was investigated by measuring its relative distribution in three intestinal regions (duodenum, ileum, and colon) and faeces using *L. plantarum* specific qPCR primers. Duodenal samples were collected 3 cm distal to the junction of the stomach and duodenum; ileal samples were collected 3 cm proximal to the junction of the ileum and cecum; and colonic samples were collected 3 cm distal to the junction of the cecum and colon. DNA was extracted from the samples using the Qiagen DNA extraction kit (Qiagen, Hilden, North Rhine-Westphalia, Germany). The sequences of *L. plantarum* specific qPCR primers are listed in Supplementary Table 2. The results are presented as mean ± S.D. normalized to the 16s rRNA expression of universal bacteria using the ΔΔCt method, in which the HFD group was used as the reference group.

### Statistical analysis

Except for the analysis of microbiota, statistical analyses were conducted using Origin2022b (OriginLab., Northampton, MA, USA). The experimental results are presented as means ± standard deviation (S.D) for 10 mice in each group and analyzed by one-way ANOVA with Tucky test for comparisons between different groups. Statistical significance was set at p value of < 0.05.

## Acknowledgements

We are grateful to Bobae Kim and Sanghun Oh for the fruitful discussions and scientific comments.

## Data availability statement

All data generated or analyzed during this study are included in this published article and its supplementary information files. Sequencing data is available using accession number PRJEB71581. All relevant data are available from the corresponding authors upon reasonable requests, after signing a data access agreement. Correspondence and requests for materials should be addressed to Ah-Ram Kim.

---

**Supplementary Figure 1.**
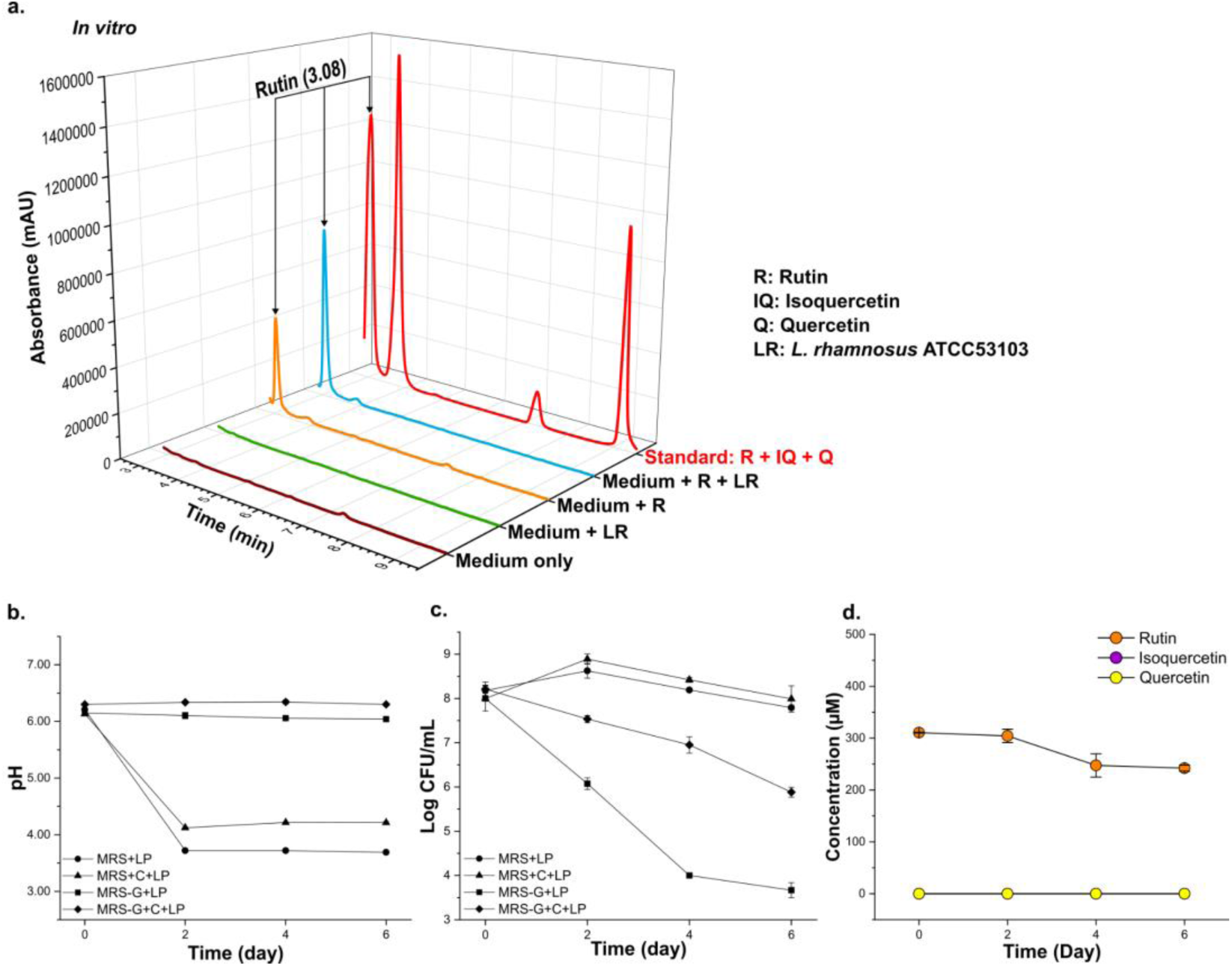
*L. rhamnosus* ATCC 53103 can not hydrolyze rutin into quercetin. (a) HPLC-DAD-MS analysis of different media associated with *L. rhamnosus* ATCC 53103 and rutin on day 6. (b-c) Among various growth media containing *L. rhmanosus* ATCC 53103, the pH exhibited the least decrease in the MRS-G+C medium, while the bacterial population was the highest in glucose-containing MRS with CaCO3 (MRS+C). (b) The pH changes of *L. rhamnosus* ATCC 53103 incubated on different mediums during the *in vitro*. (c) Growth curve of *L. rhamnosus* ATCC 53103 incubated on different mediums during the *in vitro*. (d) The concentration changes of rutin, isoquercetin and quercetin in the *L. rhamnosus* ATCC 53103 and rutin culture medium during the *in vitro*. The data are presented as mean ± SD (n = 3).

**Supplementary Figure 2.**
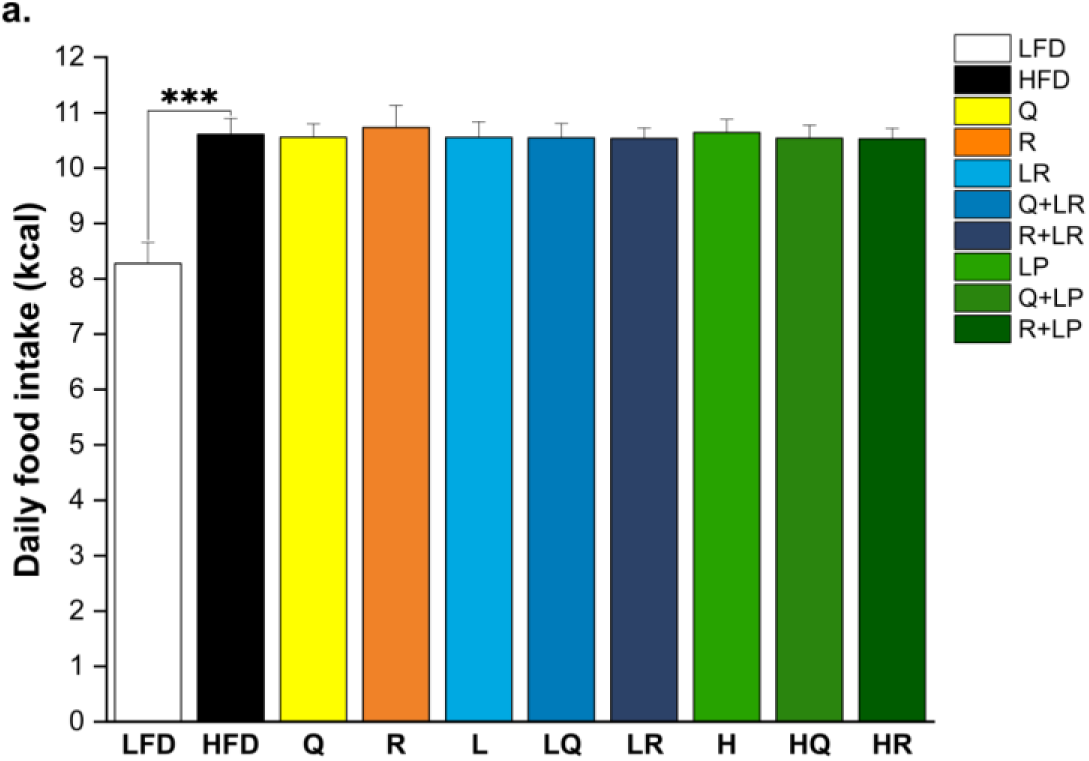
Combination of rutin and *L. plantarum* HAC03 has no effect on food intake. (a) The average daily food intake during the animal experiment. The data are presented as means ± SD (*n* = 10). One-way ANOVA with Tucky test was used for comparison with different groups. *** p < 0.001 between HFD and other groups.

**Supplementary Table 2.**
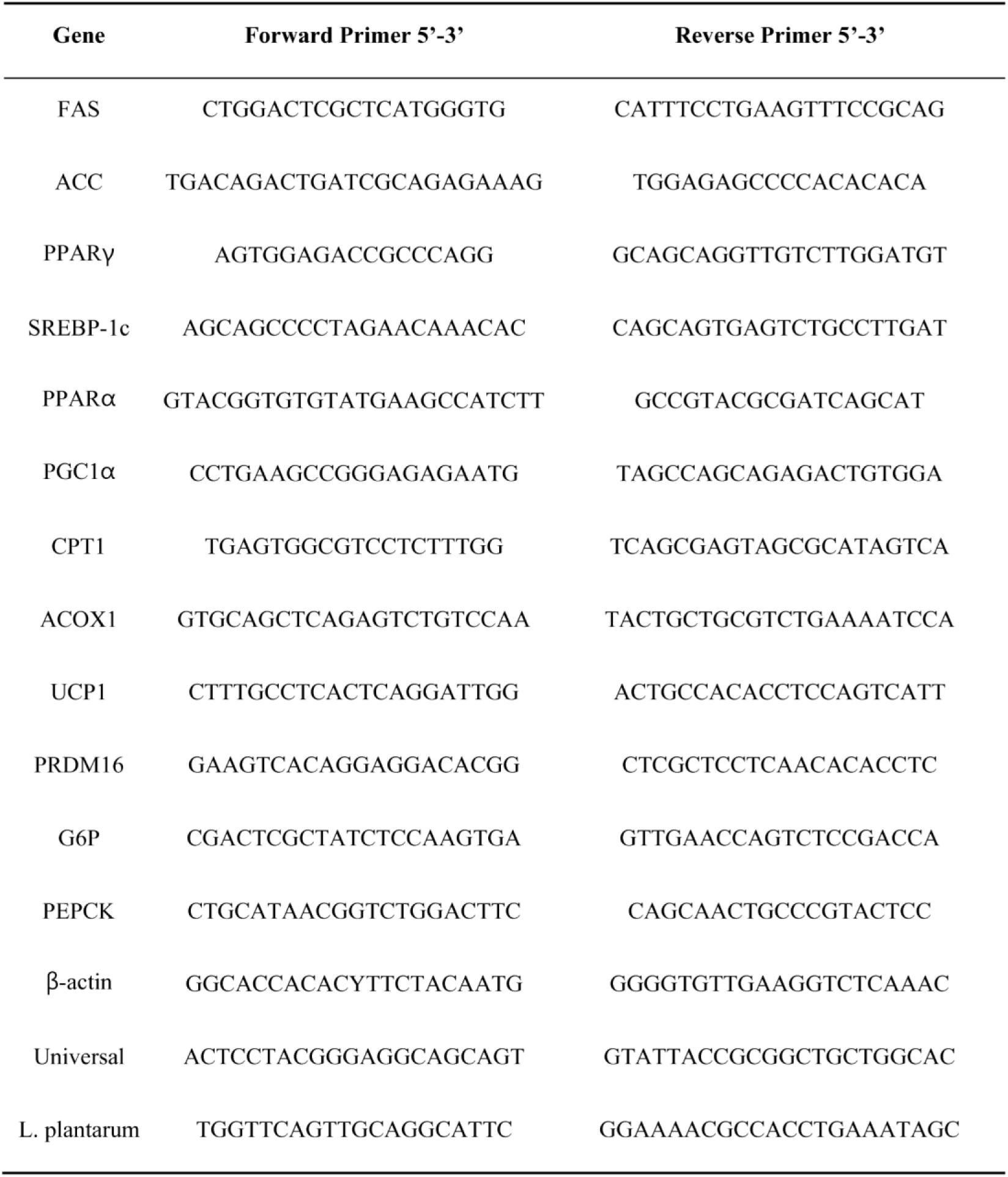
Primer sequence for qPCR.

## References

[1] Dulloo AG, Montani JP. Body composition, inflammation and thermogenesis in pathways to obesity and the metabolic syndrome: an overview. Obesity Reviews. 2012;13(S2):1–5. doi:10.1111/j.1467-789X.2012.01032.x

[2] Kyrou I, Randeva HS, Tsigos C, Kaltsas G, Weickert MO. Clinical Problems Caused by Obesity. In: Feingold KR, Anawalt B, Blackman MR, Boyce A, Chrousos G, Corpas E, et al., eds. Endotext. MDText.com, Inc.; 2000. http://www.ncbi.nlm.nih.gov/books/NBK278973/. Accessed August 30, 2023

[3] Seidell J. Obesity: a growing problem. Acta Paediatrica. 1999;88(s428):46–50. doi:10.1111/j.1651-2227.1999.tb14350.x

[4] Lobstein T, Jackson-Leach R, Powis J, Brinsden H, Gray M. World Obesity Atlas 2023. Published online March 2, 2023. https://policycommons.net/artifacts/3454894/untitled/4255209/. Accessed December 20, 2023

[5] Ivanova S, Delattre C, Karcheva-Bahchevanska D, Benbasat N, Nalbantova V, Ivanov K. Plant-Based Diet as a Strategy for Weight Control. Foods. 2021;10(12):3052. doi:10.3390/foods10123052

[6] Čižmárová B, Hubková B, Tomečková V, Birková A. Flavonoids as Promising Natural Compounds in the Prevention and Treatment of Selected Skin Diseases. International Journal of Molecular Sciences. 2023;24(7):6324. doi:10.3390/ijms24076324

[7] Yan X, Zhai Y, Zhou W, Qiao Y, Guan L, Liu H, et al. Intestinal Flora Mediates Antiobesity Effect of Rutin in High-Fat-Diet Mice. Mol Nutr Food Res. 2022;66(14):e2100948. doi:10.1002/mnfr.202100948

[8] Song D, Cheng L, Zhang X, Wu Z, Zheng X. The modulatory effect and the mechanism of flavonoids on obesity. Journal of Food Biochemistry. 2019;43(8):e12954. doi:10.1111/jfbc.12954

[9] Manach C, Morand C, Demigné C, Texier O, Régérat F, Rémésy C. Bioavailability of rutin and quercetin in rats. FEBS Letters. 1997;409(1):12–16. doi:10/dnz8zx

[10] Dias MC, Pinto DCGA, Silva AMS. Plant Flavonoids: Chemical Characteristics and Biological Activity. Molecules. 2021;26(17):5377. doi:10.3390/molecules26175377

[11] Kotik M, Kulik N, Valentová K. Flavonoids as Aglycones in Retaining Glycosidase-Catalyzed Reactions: Prospects for Green Chemistry. J Agric Food Chem. 2023;71(41):14890–14910. doi:10.1021/acs.jafc.3c04389

[12] Farias SA de S, Costa KS da, Martins J. Analysis of Conformational, Structural, Magnetic, and Electronic Properties Related to Antioxidant Activity: Revisiting Flavan, Anthocyanidin, Flavanone, Flavonol, Isoflavone, Flavone, and Flavan-3-ol. ACS omega. Published online 2021. doi:10.1021/acsomega.0c06156

[13] Xiao J. Dietary flavonoid aglycones and their glycosides: Which show better biological significance? Crit Rev Food Sci Nutr. 2017;57(9):1874–1905. doi:10/gmxg2b

[14] Butun B, Topcu G, Ozturk T. Recent Advances on 3-Hydroxyflavone Derivatives: Structures and Properties. Mini-Reviews in Medicinal Chemistry. 18(2):98–103.

[15] Křen V. Glycoside vs. Aglycon: The Role of Glycosidic Residue in Biological Activity. In: Fraser-Reid BO, Tatsuta K, Thiem J, eds. Glycoscience: Chemistry and Chemical Biology. Springer; 2008:2589–2644. doi:10.1007/978-3-540-30429-6_67

[16] Hai Y, Zhang Y, Liang Y, Ma X, Qi X, Xiao J, et al. Advance on the absorption, metabolism, and efficacy exertion of quercetin and its important derivatives. Food Frontiers. 2020;1(4):420–434. doi:10/gmtn4d

[17] Zhang H, Hassan YI, Liu R, Mats L, Yang C, Liu C, et al. Molecular Mechanisms Underlying the Absorption of Aglycone and Glycosidic Flavonoids in a Caco-2 BBe1 Cell Model. ACS Omega. 2020;5(19):10782–10793. doi:10.1021/acsomega.0c00379

[18] AL-Ishaq RK, Liskova A, Kubatka P, Büsselberg D. Enzymatic Metabolism of Flavonoids by Gut Microbiota and Its Impact on Gastrointestinal Cancer. Cancers (Basel). 2021;13(16):3934. doi:10.3390/cancers13163934

[19] Braune A, Blaut M. Bacterial species involved in the conversion of dietary flavonoids in the human gut. Gut Microbes. 2016;7(3):216–234. doi:10/gjvhtt

[20] Beekwilder J, Marcozzi D, Vecchi S, de Vos R, Janssen P, Francke C, et al. Characterization of Rhamnosidases from Lactobacillus plantarum and Lactobacillus acidophilus. Appl Environ Microbiol. 2009;75(11):3447–3454. doi:10.1128/AEM.02675-08

[21] Michlmayr H, Kneifel W. β-Glucosidase activities of lactic acid bacteria: mechanisms, impact on fermented food and human health. FEMS Microbiology Letters. 2014;352(1):1–10. doi:10.1111/1574-6968.12348

[22] Valentová K, Vrba J, Bancířová M, Ulrichová J, Křen V. Isoquercitrin: Pharmacology, toxicology, and metabolism. Food and Chemical Toxicology. 2014;68:267–282. doi:10.1016/j.fct.2014.03.018

[23] Hollman PCH. Absorption, Bioavailability, and Metabolism of Flavonoids. Pharmaceutical Biology. 2004;42(sup1):74–83. doi:10.3109/13880200490893492

[24] Jiang H, Horiuchi Y, Hironao K yu, Kitakaze T, Yamashita Y, Ashida H. Prevention effect of quercetin and its glycosides on obesity and hyperglycemia through activating AMPKα in high-fat diet-fed ICR mice. Journal of Clinical Biochemistry and Nutrition. 2020;67(1):74. doi:10.3164/jcbn.20-47

[25] Mueller M, Zartl B, Schleritzko A, Stenzl M, Viernstein H, Unger FM. Rhamnosidase activity of selected probiotics and their ability to hydrolyse flavonoid rhamnoglucosides. Bioprocess Biosyst Eng. 2018;41(2):221–228. doi:10.1007/s00449-017-1860-5

[26] Salek S S., van Turnhout A G., Kleerebezem R, van Loosdrecht M C. M. pH control in biological systems using calcium carbonate. Biotechnology and Bioengineering. 2015;112(5):905–913. doi:10.1002/bit.25506

[27] McMahon H, Zoecklein BW, Fugelsang K, Jasinski Y. Quantification of glycosidase activities in selected yeasts and lactic acid bacteria. J Ind Microbiol Biotech. 1999;23(3):198–203. doi:10.1038/sj.jim.2900720

[28] Zhang Z, Zhou Z, Li Y, Zhou L, Ding Q, Xu L. Isolated exopolysaccharides from Lactobacillus rhamnosus GG alleviated adipogenesis mediated by TLR2 in mice. Sci Rep. 2016;6(1):36083. doi:10.1038/srep36083

[29] Gorbach S, Doron S, Magro F. Chapter 7 - Lactobacillus rhamnosus GG. In: Floch MH, Ringel Y, Allan Walker W, eds. The Microbiota in Gastrointestinal Pathophysiology. Academic Press; 2017:79–88. doi:10.1016/B978-0-12-804024-9.00007-0

[30] Cui XB, Chen SY. White adipose tissue browning and obesity. J Biomed Res. 2017;31(1):1–2. doi:10.7555/JBR.31.20160101

[31] Kahn BB, Flier JS. Obesity and insulin resistance. J Clin Invest. 2000;106(4):473℃481. doi:10.1172/JCI10842

[32] Polyzos SA, Kountouras J, Mantzoros CS. Obesity and nonalcoholic fatty liver disease: From pathophysiology to therapeutics. Metabolism. 2019;92:82–97. doi:10.1016/j.metabol.2018.11.014

[33] Ahmad A, Yang W, Chen G, Shafiq M, Javed S, Ali Zaidi SS, et al. Analysis of gut microbiota of obese individuals with type 2 diabetes and healthy individuals. PLoS One. 2019;14(12):e0226372. doi:10.1371/journal.pone.0226372

[34] Lippert K, Kedenko L, Antonielli L, Kedenko I, Gemeier C, Leitner M, et al. Gut microbiota dysbiosis associated with glucose metabolism disorders and the metabolic syndrome in older adults. Benef Microbes. 2017;8(4):545–556. doi:10.3920/BM2016.0184

[35] Dupuit M, Chavanelle V, Chassaing B, Perriere F, Etienne M, Plissonneau C, et al. The TOTUM-63 Supplement and High-Intensity Interval Training Combination Limits Weight Gain, Improves Glycemic Control, and Influences the Composition of Gut Mucosa-Associated Bacteria in Rats on a High Fat Diet. Nutrients. 2021;13(5):1569. doi:10.3390/nu13051569

[36] Islam MdR, Hassan YI, Das Q, Lepp D, Hernandez M, Godfrey DV, et al. Dietary organic cranberry pomace influences multiple blood biochemical parameters and cecal microbiota in pasture-raised broiler chickens. Journal of Functional Foods. 2020;72:104053. doi:10.1016/j.jff.2020.104053

[37] Queipo-Ortuño MI, Boto-Ordóñez M, Murri M, Gomez-Zumaquero JM, Clemente-Postigo M, Estruch R, et al. Influence of red wine polyphenols and ethanol on the gut microbiota ecology and biochemical biomarkers. Am J Clin Nutr. 2012;95(6):1323–1334. doi:10.3945/ajcn.111.027847

[38] Muvhulawa N, Dludla PV, Ziqubu K, Mthembu SXH, Mthiyane F, Nkambule BB, et al. Rutin ameliorates inflammation and improves metabolic function: A comprehensive analysis of scientific literature. Pharmacological Research. 2022;178:106163. doi:10.1016/j.phrs.2022.106163

[39] Ganeshpurkar A, Saluja AK. The Pharmacological Potential of Rutin. Saudi Pharmaceutical Journal. 2017;25(2):149–164. doi:10.1016/j.jsps.2016.04.025

[40] Andres S, Pevny S, Ziegenhagen R, Bakhiya N, Schäfer B, Hirsch-Ernst KI, et al. Safety Aspects of the Use of Quercetin as a Dietary Supplement. Molecular Nutrition & Food Research. 2018;62(1):1700447. doi:10/ghwxnn

[41] Okamoto T. Safety of quercetin for clinical application (Review). International Journal of Molecular Medicine. 2005;16(2):275–278. doi:10.3892/ijmm.16.2.275

[42] Panche AN, Diwan AD, Chandra SR. Flavonoids: an overview. J Nutr Sci. 2016;5:e47. doi:10.1017/jns.2016.41

[43] Pei R, Liu X, Bolling B. Flavonoids and gut health. Current Opinion in Biotechnology. 2020;61:153–159. doi:10/gmjrtc

[44] Varshney R, Mishra R, Das N, Sircar D, Roy P. A comparative analysis of various flavonoids in the regulation of obesity and diabetes: An in vitro and in vivo study. Journal of Functional Foods. 2019;59:194–205. doi:10.1016/j.jff.2019.05.004

[45] Yang J, Lee J, Kim Y. Effect of Deglycosylated Rutin by Acid Hydrolysis on Obesity and Hyperlipidemia in High-Fat Diet-Induced Obese Mice. Nutrients. 2020;12(5):E1539. doi:10.3390/nu12051539

[46] Semwal R, Joshi SK, Semwal RB, Semwal DK. Health benefits and limitations of rutin - A natural flavonoid with high nutraceutical value. Phytochemistry Letters. 2021;46:119–128. doi:10.1016/j.phytol.2021.10.006

[47] Uskova MA, Kravchenko LV, Avrenjeva LI, Tutelyan VA. Effect of Lactobacillus casei 114001 Probiotic on Bioactivity of Rutin. Bull Exp Biol Med. 2010;149(5):578–582. doi:10.1007/s10517-010-0997-x

[48] Mazzeo MF, Lippolis R, Sorrentino A, Liberti S, Fragnito F, Siciliano RA. Lactobacillus acidophilus—Rutin Interplay Investigated by Proteomics. PLoS One. 2015;10(11):e0142376. doi:10.1371/journal.pone.0142376

[49] Park CM, Kim GM, Cha GS. Biotransformation of Flavonoids by Newly Isolated and Characterized Lactobacillus pentosus NGI01 Strain from Kimchi. Microorganisms. 2021;9(5):1075. doi:10/gmxgz9

[50] Erlund I, Kosonen T, Alfthan G, Mäenpää J, Perttunen K, Kenraali J, et al. Pharmacokinetics of quercetin from quercetin aglycone and rutin in healthy volunteers. Eur J Clin Pharmacol. 2000;56(8):545–553. doi:10.1007/s002280000197

[51] Riva A, Kolimár D, Spittler A, Wisgrill L, Herbold CW, Abrankó L, et al. Conversion of Rutin, a Prevalent Dietary Flavonol, by the Human Gut Microbiota. Front Microbiol. 2020;11:585428. doi:10.3389/fmicb.2020.585428

[52] Chen Z, Zheng S, Li L, Jiang H. Metabolism of flavonoids in human: a comprehensive review. Curr Drug Metab. 2014;15(1):48–61. doi:10.2174/138920021501140218125020

[53] Kolb H. Obese visceral fat tissue inflammation: from protective to detrimental? BMC Medicine. 2022;20(1):494. doi:10.1186/s12916-022-02672-y

[54] Gu S, Chen D, Zhang JN, Lv X, Wang K, Duan LP, et al. Bacterial community mapping of the mouse gastrointestinal tract. PLoS One. 2013;8(10):e74957. doi:10.1371/journal.pone.0074957

[55] Anders JL, Moustafa MAM, Mohamed WMA, Hayakawa T, Nakao R, Koizumi I. Comparing the gut microbiome along the gastrointestinal tract of three sympatric species of wild rodents. Sci Rep. 2021;11(1):19929. doi:10.1038/s41598-021-99379-6

[56] Mentula S, Harmoinen J, Heikkilä M, Westermarck E, Rautio M, Huovinen P, et al. Comparison between cultured small-intestinal and fecal microbiotas in beagle dogs. Appl Environ Microbiol. 2005;71(8):4169–4175. doi:10.1128/AEM.71.8.4169-4175.2005

[57] Park S, Ji Y, Park H, Lee K, Park H, Beck BR, et al. Evaluation of functional properties of lactobacilli isolated from Korean white kimchi. Food Control. 2016;69:5–12. doi:10.1016/j.foodcont.2016.04.037

[58] King WL, Siboni N, Williams NLR, Kahlke T, Nguyen KV, Jenkins C, et al. Variability in the Composition of Pacific Oyster Microbiomes Across Oyster Families Exhibiting Different Levels of Susceptibility to OsHV-1 μvar Disease. Frontiers in Microbiology. 2019;10. https://www.frontiersin.org/articles/10.3389/fmicb.2019.00473. Accessed January 3, 2023

[59] Callahan BJ, McMurdie PJ, Rosen MJ, Han AW, Johnson AJA, Holmes SP. DADA2: High-resolution sample inference from Illumina amplicon data. Nat Methods. 2016;13(7):581–583. doi:10.1038/nmeth.3869

[60] Glöckner FO, Yilmaz P, Quast C, Gerken J, Beccati A, Ciuprina A, et al. 25 years of serving the community with ribosomal RNA gene reference databases and tools. J Biotechnol. 2017;261:169–176. doi:10.1016/j.jbiotec.2017.06.1198

[61] Bolyen E, Rideout JR, Dillon MR, Bokulich NA, Abnet CC, Al-Ghalith GA, et al. Reproducible, interactive, scalable and extensible microbiome data science using QIIME 2. Nat Biotechnol. 2019;37(8):852–857. doi:10.1038/s41587-019-0209-9

[62] Lahti L, Shetty S. microbiome R package. Published online 2017. doi:10.18129/B9.bioc.microbiome

