## Supplementary figures and images for "Inducing a synergistic anti-obesity effect by increasing the bioavailability of the flavonoid rutin with a *L. plantarum* species"

### supplementary_table2

#### Supplementary Table 2. Primer sequence for qPCR

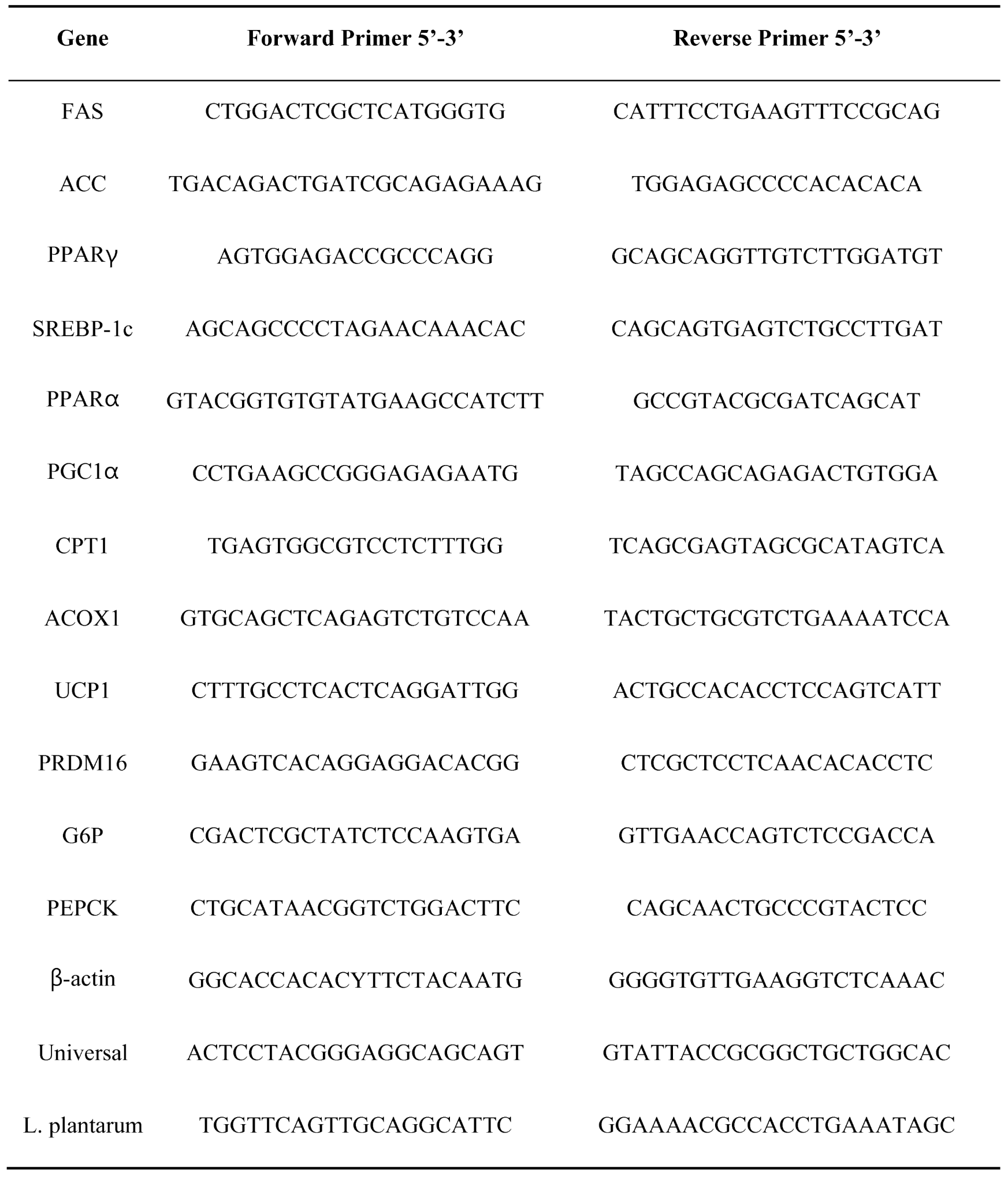
