## supplementary_figure2 for "Inducing a synergistic anti-obesity effect by increasing the bioavailability of the flavonoid rutin with a *L. plantarum* species"

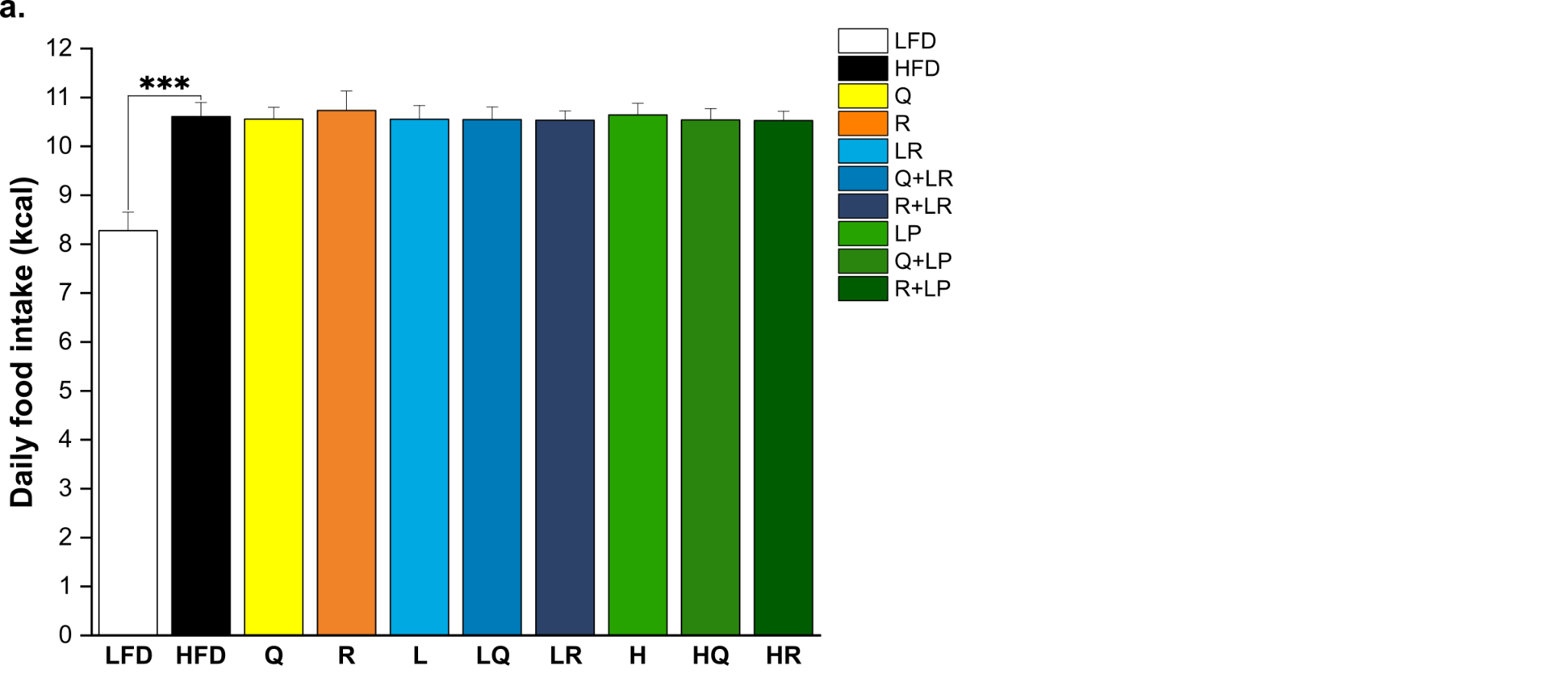

**Supplementary Figure 2. Combination of rutin and *L. plantarum* HAC03 has no effect on food intake**. (a) The average daily food intake during the animal experiment. The data are presented as means ± SD (*n* = 10). One-way ANOVA with Tucky test was used for comparison with different groups. *** p < 0.001 between HFD and other groups.
