## supplementary_figure1 for "Inducing a synergistic anti-obesity effect by increasing the bioavailability of the flavonoid rutin with a *L. plantarum* species"

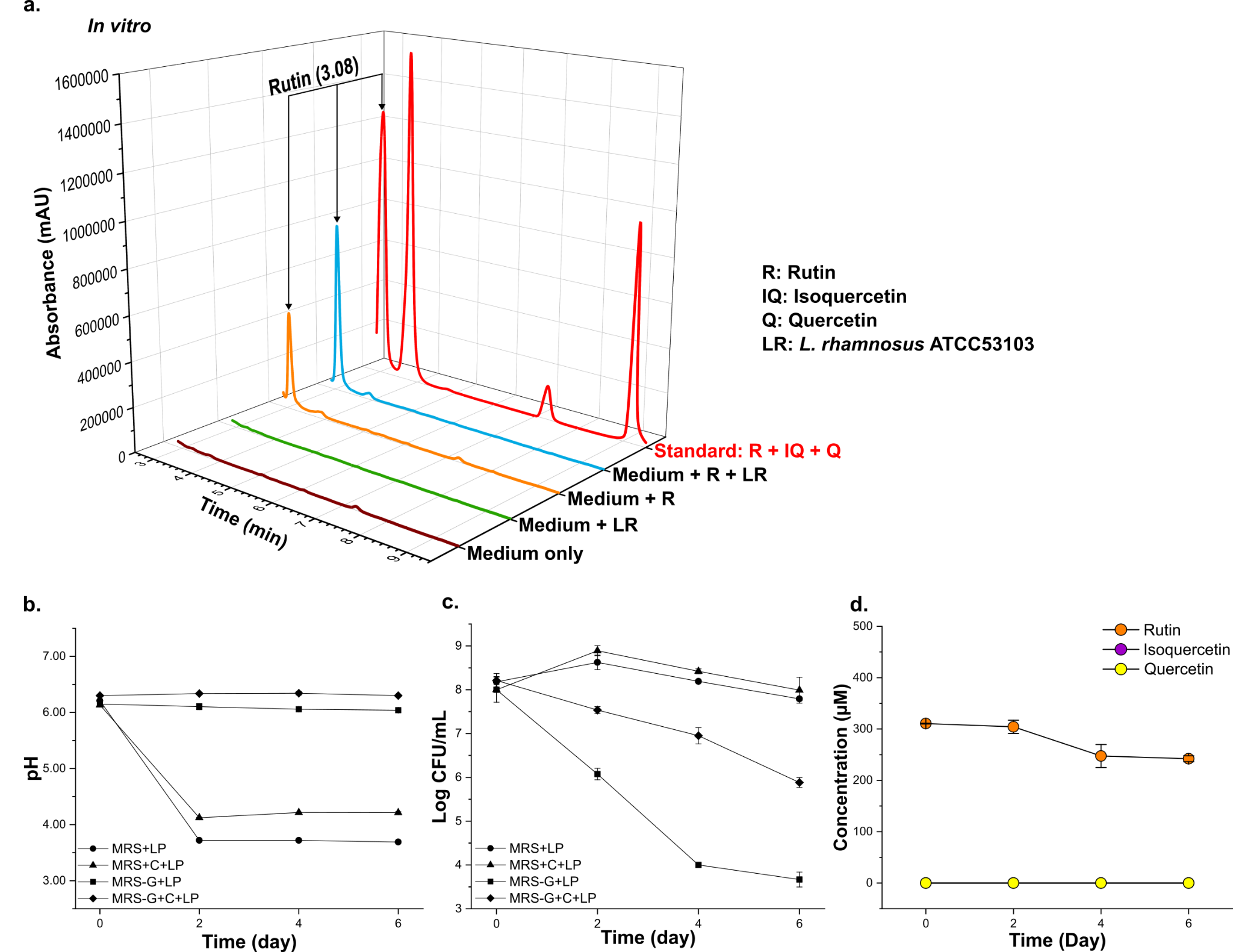

### Supplementary Figure 1. *L. rhamnosus* ATCC 53103 can not hydrolyze rutin into quercetin. (a) HPLC-DAD-MS analysis of different media associated with *L. rhamnosus* ATCC 53103 and rutin on day 6. (b-c) Among various growth media containing *L. rhmanosus* ATCC 53103, the pH exhibited the least decrease in the MRS-G+C medium, while the bacterial population was the highest in glucose-containing MRS with CaCO3 (MRS+C). (b) The pH changes of *L. rhamnosus* ATCC 53103 incubated on different mediums during the *in vitro*. (c) Growth curve of *L. rhamnosus* ATCC 53103 incubated on different mediums during the *in vitro*. (d) The concentration changes of rutin, isoquercetin and quercetin in the *L. rhamnosus* ATCC 53103 and rutin culture medium during the *in vitro*. The data are presented as mean ± SD (n = 3).
